# Dissecting Functional, Structural, and Molecular Requirements for Serotonin Release from Mouse Enterochromaffin Cells

**DOI:** 10.1101/2021.05.28.446100

**Authors:** Ahmed Shaaban, Frederike Maaß, Valentin Schwarze, Mari L. Lund, Sabine Beuermann, Michelle Chan, Christiane Harenberg, Gavin A. Bewick, Damien J. Keating, Fritz Benseler, Benjamin H. Cooper, Cordelia Imig

## Abstract

Serotonergic enterochromaffin (EC) cells of the gut epithelium are secretory sensory cells that communicate with vagal neurons. EC cells exhibit many features of neurons in the brain, raising the hypothesis that synapse-like contacts may mediate fast and directed signalling. To dissect functional, structural, and molecular properties underlying serotonin release from genetically identified EC cells, we employed a multidisciplinary *in vitro* approach combining intestinal epithelial cell and organoid cultures, electrochemistry, correlated light- and electron microscopy, and gene expression and biochemical analyses. Despite the presence of key molecules of the synaptic neurotransmitter release machinery, we found that the majority of serotonin is released with slow kinetics from large dense-core rather than small synaptic-like vesicles. While we cannot exclude synapse-like transmission between EC cells and neurons *in vivo*, our data support the notion that the predominant mode of serotonin secretion is similar to that of other endocrine cell types.

## Introduction

The intestinal epithelium serves as an interface between the body and the environment. As such, it carries out important functions that include food digestion, nutrient absorption, and the detection of changes in the gut milieu^1^. Information exchange between the gut and the brain is particularly critical for the regulation of physiology, metabolism, and behaviours, including the regulation of appetite and satiety and consequential food intake^2^. An important subpopulation of cells in this context are enteroendocrine cells (EECs). EECs are secretory cells that only constitute ∼1% of all gut epithelial cells^3^ and are found scattered at low spatial density throughout the entire gastrointestinal (GI) tract^4^. EECs are chemo- and mechanosensitive and respond to the activation of molecular receptors by mechanical force^5, 6^, nutrients, metabolites, irritants, and toxins^7, 8^ via the release of a variety of different peptide hormones (e.g. cholecystokinin, somatostatin, proglucagon, glucose-dependent insulinotropic polypeptide, neurotensin, secretin, and substance P^9–11^) and neurotransmitters (i.e. glutamate^12^, serotonin^13^, and ATP^14^). Different secretory products are often found co-expressed within the same cell thereby rendering EECs a highly diverse group of cells^1, 11, 15^. Interestingly, some EEC subtypes also express transcripts that encode proteins involved in neurotransmitter release from neuronal synapses in the brain^12, 16, 17^ and possess a basolateral process termed ‘neuropod’^12, 18, 19^. It is therefore generally believed that at least some EECs form synapses with vagal sensory afferents and that this signalling route may be critically important in mediating rapid (hundreds of milliseconds) gut-brain signalling^12, 19^. Despite this knowledge, it is still largely unknown how such synapse-like contact points are organized at the ultrastructural level, which molecules mediate the fusion of secretory vesicles in response to Ca^2+^, or what the kinetics of release are for signalling molecules in different EEC subtypes. This knowledge gap can be largely attributed to the fact that the low-density distribution of EECs in the GI tract has long posed severe technical difficulties for high-resolution biochemical, morphological, or functional analyses. Determining the kinetics of vesicle fusion events, the amount of release from individual vesicles and cells, and the structural and molecular organization of this process will, however, contribute towards a better understanding of how information is sensed in the gut and relayed by EECs to the brain.

In the present study, we set up a multidisciplinary experimental approach to systematically explore EEC structure, function, and cell biology *in vitro*. We focused our efforts on serotonergic enterochromaffin (EC) cells that comprise the largest EEC subgroup. EC cells are found in all regions of the GI tract^4^, and they store together the majority of the body’s serotonin (5-hydroxytryptamine, 5-HT)^20–22^. EC cells are electrically excitable, however action potential firing is not essential for P/Q-type volage-gated calcium channel-mediated Ca^2+^-triggering of exocytosis^17, 23^. Gut-derived 5-HT mediates diverse body functions such as the regulation of gut motility^21, 22^ and serves as a fast-acting satiety signal^24, 25^. Consequently, altered 5-HT release from EC cells has been associated with a variety of different disorders, including obesity, inflammatory bowel diseases, irritable bowel syndrome, as well as visceral hypersensitivity, nausea, and vomiting^26, 27^. The rate-limiting enzyme of the 5-HT synthesis in EC cells is the non-neuronal tryptophan hydroxylase 1 (Tph1)^28^ and Tph1 expression serves as a highly reliable molecular marker to discriminate EC cells from other EEC subtypes^15, 29–31^. EC cells themselves, however, form a heterogeneous cell population^32^ in that they change their molecular identity during their lifetime, which is marked by a shift in co-expression from substance P to secretin during maturation^30, 31, 33^ and co-expression of 5-HT with CCK, PYY, and GLP-1 has also been reported^32^. Finally, EC cells in the small and large intestine exhibit differences in their morphologies^34, 35^ and in their repertoire of expressed nutrient- and metabolite receptors^36–38^. The close proximity of EC cells to ionotropic 5-HT3 receptor-expressing vagal sensory afferents^17^ and the previously described extremely fast 5-HT release kinetics from acutely isolated EC cells^39, 40^ have made them prime candidates to carry out fast and directed information transfer from the gut to the brain. Despite the physiological importance of EC cells and a growing understanding of the molecular mechanisms by which they detect different sensory modalities in the gut lumen^5, 6, 17, 23^, very little is known about how excitation-secretion coupling is achieved in EC cells.

## Results

### An experimental workflow for the study of EC cells on the single cell level

To determine functional, structural, and molecular requirements for 5-HT release from mouse EC cells, we setup an experimental workflow combining 2D and 3D mouse intestinal epithelial cell cultures, single-cell electrochemistry, and correlative light and electron microscopy (CLEM) (Fig.1).

**Figure 1.**
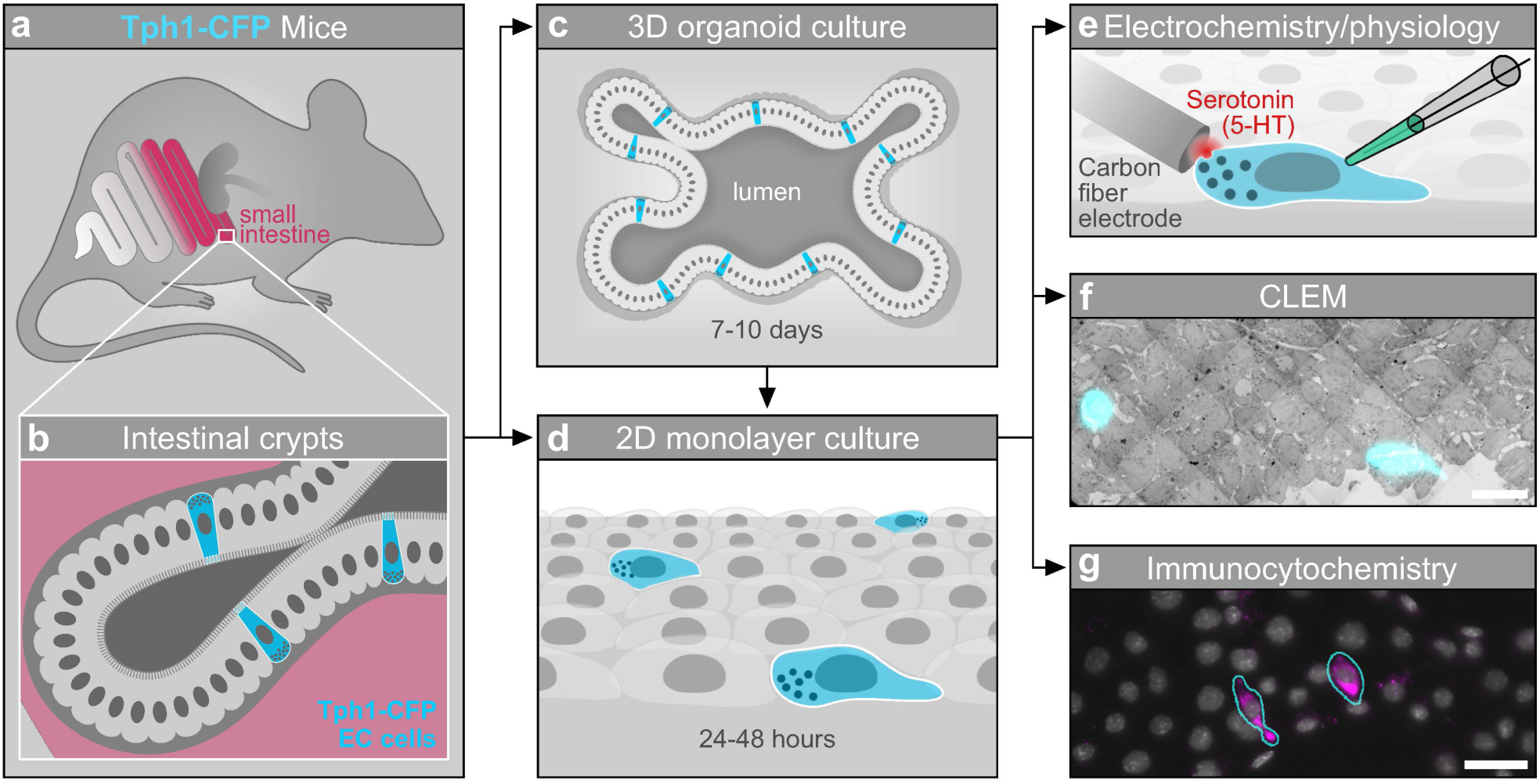
Experimental Workflow for Studying Functional, Structural, and Molecular Aspects of Mouse Enterochromaffin (EC) Cells in Culture. **a** Transgenic Tph1-CFP mice express the fluorescent reporter cyan fluorescent protein (CFP) in EC cells of the gut epithelium throughout the gastrointestinal tract. **b** Stem cell-containing intestinal crypts isolated from the proximal small intestine of Tph1-CFP mice and taken into culture generate 3D intestinal organoids (**c**) or acute 2D epithelial monolayers (**d**). **c**-**d** Mature organoids (**c**) may be dissociated and plated as 2D monolayers (**d**) for subsequent experiments. **e** CFP-positive EC cells are accessible for electrophysiology and electrochemistry to monitor 5-HT secretion from individual vesicles with high temporal resolution. **f** Correlated light- and electron microscopy (CLEM) enables ultrastructural studies to determine the organization of secretory granules in EC cells. **g** Immunofluorescent light microscopic imaging of cultured EC cells to resolve the subcellular distribution of candidate molecular components of the secretory machinery. Scale bars: 20 *µ*m (**f** and **g**)

To achieve this, we took advantage of the fact that the intestinal epithelium is a rapidly renewing tissue that replaces its cellular components continuously from adult intestinal stem cells that reside in the crypts. These crypts can be isolated from gut tissue of transgenic fluorescent reporter mice for distinct EEC types including a transgenic mouse line that expresses cyan fluorescent protein (CFP) under the control of the mouse tryptophan hydroxylase 1 (Tph1) gene (Fig. 1a and b) and therefore specifically in EC cells^6, 23, 29^. Crypts are then taken into culture to form mature 3D organoids in a hydrogel environment within 1-2 weeks (Fig. 1c). This so-called ‘mini gut’ *in vitro* tissue model system allows the recapitulation of many physiological and morphological features of the *in vivo* epithelium in a culture setting (i.e. polarized epithelial layer containing all cell types including all distinct EEC types^41^). Organoids grown in a hydrogel environment can be kept like a cell line in culture for many months, and stem cells that reside within the organoids continue to proliferate and can be manipulated genetically to generate mutant organoid cell lines^42^. In addition to the 3D organoid system, primary acute 2D epithelial monolayer cultures can be prepared from freshly isolated crypts (Fig. 1d), which have the advantage that cells are more easily accessible for high-resolution functional and imaging experiments^43, 44^. Finally, mature organoids can be removed from Matrigel and converted into planar 2D-epithelial monolayer cultures for light microscopy, electron microscopy, as well as electrophysiology and electrochemistry experiments^45^ (Fig. 1e-g). Parts of this study were previously presented^46, 47^.

### Validation of the Tph1-CFP mouse line

We first set out to characterise the expression of the CFP fluorescent reporter in intestinal cells of Tph1-CFP animals in different preparations (Fig. 2). Live imaging experiments consistently revealed the presence of strong CFP fluorescence in individual cells in both, duodenal monolayer cultures (Fig. 2a-c) and organoids (Fig. 2d-f). Moreover, strong endogenous CFP fluorescence was also retained after immersion fixation of duodenal (Fig. 2g-i) and colonic (Fig. 2j-l) tissue fragments in paraformaldehyde-based fixatives.

**Figure 2.**
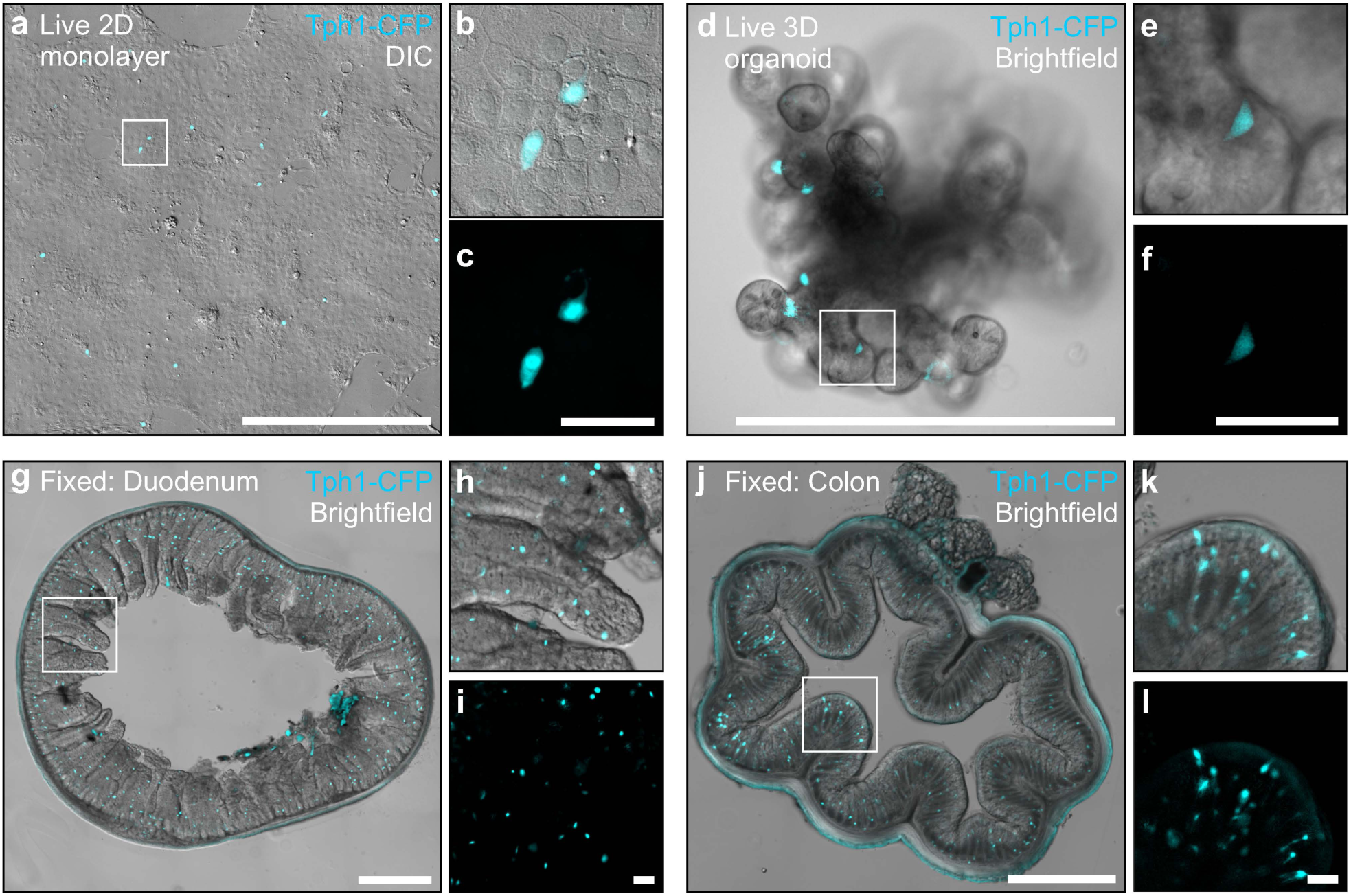
Characterization of Endogenous CFP Fluorescence in Cells of the Intestinal Epithelium of Tph1-CFP Mice in Different Preparations. **a**-**f** Live imaging of cultures derived from Tph1-CFP animals indicate strong endogenous CFP fluorescence in epithelial monolayers (**a**-**c**) and 3D organoids (**d**-**f**). **g**-**l** Endogenous CFP fluorescence is preserved in aldehyde-fixed material of the proximal small intestine (**g**-**i**) and colon (**j**-**l**). Scale bars: 500 *µ*m (**a**, **d**, **g**, and **j**); 50 *µ*m (**c**, **f**, **i**, and **l**).

We then explored whether all Tph1-CFP positive cells were also 5-HT-positive and vice versa, whether all 5-HT-positive cells expressed the fluorescent reporter. For this we performed immunocytochemistry on duodenal 2D epithelial monolayer cultures from Tph1-CFP mice using antibodies directed against 5-HT and green fluorescent protein (GFP) (Supplementary Fig. S1a-k). The latter antibody also reacts with CFP and can therefore be used to amplify the endogenous fluorescence of the reporter to guarantee that also cells with a weak endogenous CFP fluorescence were included in the analysis. We found that 100% of anti-GFP-positive cells also contain CFP and likewise 100% 5-HT (Supplementary Fig. S1i) and that 92.62 ± 8% and 91.31 ± 10.3% (mean ± SD) of all serotonergic cells exhibited an endogenous CFP or an anti-GFP amplified signal, respectively (Supplementary Fig. S1k). Throughout our experiments, we noticed apparent differences in the intensity of the endogenous CFP-signal between individual fluorescent cells. To test whether differences in the degree of maturity of EC cells could explain this finding, we stained Tph1-CFP monolayer cultures with antibodies directed against secretin and substance P (Supplementary Fig. S1l-s). Both peptides are known to be co-expressed with serotonin in mouse EC cells, however, they are rarely found together in the same cell^33^. Instead, they delimit distinct EC cell populations of different maturity within the intestinal epithelium^30, 31, 33^. In line with previous studies, qualitative co-localization analysis of CFP-positive cells revealed that cells with weak endogenous CFP-fluorescence were preferentially substance P-positive, whereas strong CFP-fluorescence correlated with secretin-immunoreactivity.

In summary, we demonstrate that CFP-expression in duodenal monolayer cultures from Tph1-CFP animals is a highly specific fluorescent marker for 5-HT-positive EC cells, and that strongly fluorescent cells likely correspond to more mature EC cells that co-express preferentially secretin^30, 31, 33^.

### Electrochemical detection of 5-HT released from single EC cell Vesicles

We next combined carbon-fibre amperometry and whole-cell patch clamp electrophysiology on EC cells in acutely generated duodenal monolayer cultures to measure the amount and kinetics of 5-HT secretion from EC cells with single-vesicle resolution and at real-time (Fig. 3a-p). In these experiments, we approached an individual CFP-positive EC cell with a 5 *µ*m diameter carbon fibre electrode held at a positive potential (+650 mV) for the electrochemical detection of oxidized monoamines^48, 49^ (Fig. 3a). The cell was simultaneously infused with a solution with a [Ca^2+^] < 5 *µ*m through the patch pipette to trigger the fusion of secretory vesicles with the plasma membrane^50^. This concentration is considered “moderate” for other neuroendocrine cell types^51^. For the analysis of amperometric recordings (Fig. 3b), we used a spike detection threshold of 5 pA. This criterion was chosen to make sure that current transients clearly exceeded the average baseline noise in all recordings since we did not experimentally determine the noise levels for each cell measured individually. Each current “spike” corresponds to the release of 5-HT from a single exocytotic vesicle fusion event^48^.

**Figure 3.**
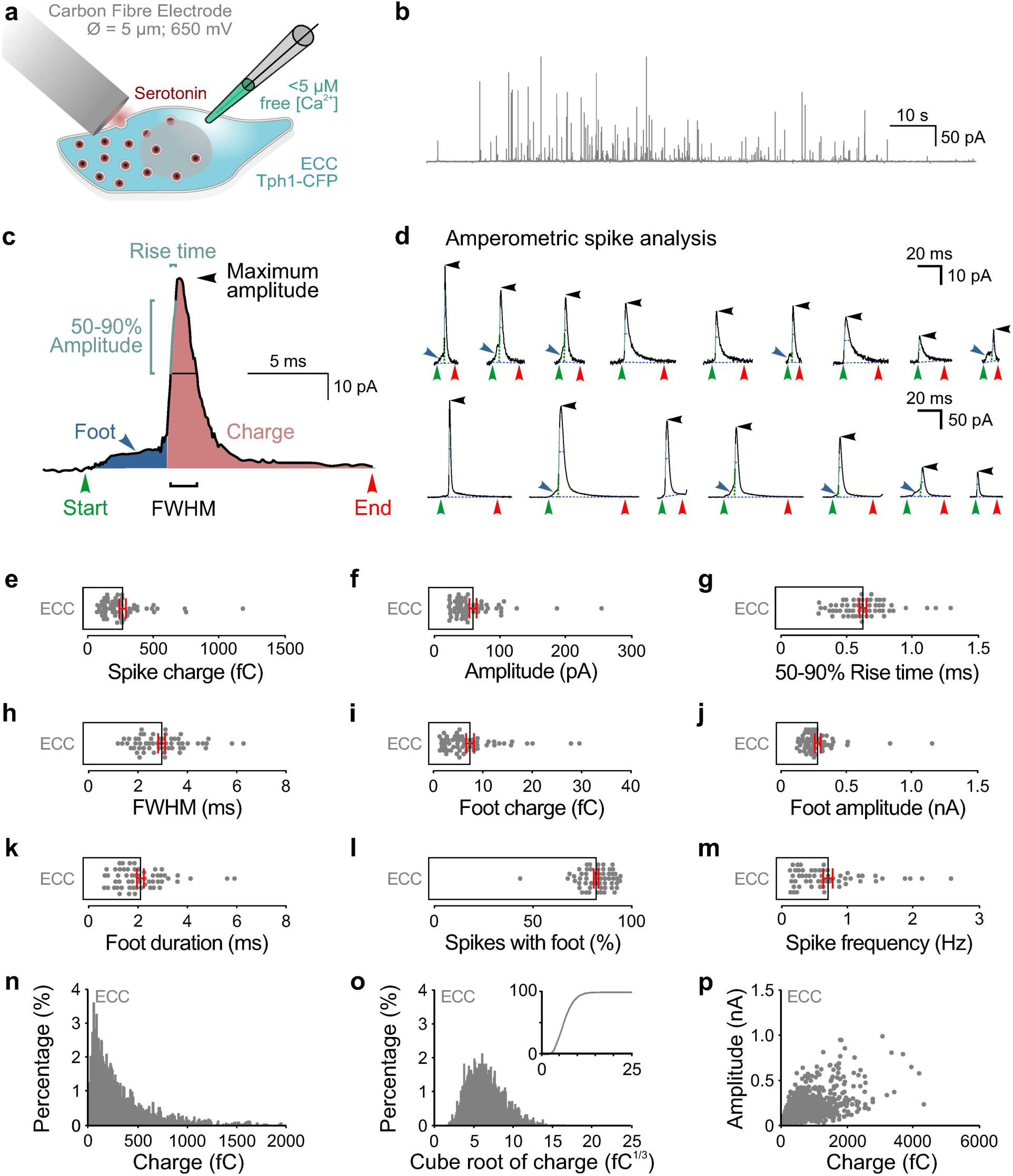
Single Cell-Carbon Fiber Amperometry for the Direct Detection of 5-HT released from EC Cells in Epithelial Monolayers. **a** Illustration of the experimental design. Exocytosis was triggered by infusing individual CFP-positive EC cells with a solution containing moderate (< 5 *µ*m) [Ca^2+^]. Serotonin released from single fusing vesicles was oxidized at the carbon-fiber electrode held at +650 mV. Currents generated by electrons transferred in the redox reaction were recorded as amperometric “spikes” for 120 s after opening the cell. **b** Representative amperometric recording of a Tph1-CFP EC cell. **c** Illustration of an individual amperometric spike and of the parameters analyzed. **d** Examples of amperometric currents evoked by the oxidation of 5-HT with variable kinetics. Note the difference in the scale between the upper and lower panels. **e** Median spike charge (fC). **f** Median maximum spike amplitude (pA). **g** Median 50-90% rise time (ms). **h** Median full-width half maximum (FWHM) corresponding to the width (ms) at half maximal amplitude. **i** Median foot charge (fC). **j** Median maximum foot amplitude (pA). **k** Median foot duration (ms). **l** Percentage of spikes with a foot signal from all spikes analyzed per cell (%). **m** Average spike frequency (Hz) per cell recorded during the 120 s recording interval before excluding overlapping spikes for the analysis of spike properties (**d**-**k**). **n** Frequency distribution of the amperometric spike charge (fC) for all events measured over all cells [bin width 10 fC; only events <2000 pA; 3216 spikes]. **o** Frequency distribution plotting the cube-root of amperometric spike charges (fC^1/3^) for all events measured over all cells [bin width 0.1 fC^1/3^; 3254 spikes]. Insert, cumulative frequency distribution for the same data. ***p*** Scatterplot illustrating the relationship between amperometric spike charge (fC) and maximum amplitude (nA) for each event measured [3254 spikes]. Error bars indicate mean ± SEM; 6 cultures, 57 cells.

Quantifications of spike parameters (Fig. 3c and d) such as the spike charge and maximal current amplitude (Fig. 3e and f) permit an estimation about the amount of transmitter released from a single vesicle. It has previously been shown by carbon fibre amperometry that the amount of 5-HT released from individual vesicles in serotonergic Retzius neurons of the leech correlate well with size of the vesicles as assessed by electron microscopy^49^. These data provided support for the notion that the concentration of 5-HT in vesicles of different sizes is constant and they imply that the charge of an amperometric current may serve as a reliable proxy for the volume of a secretory vesicle. We found here that the charge distribution for EC cells was highly skewed with an average median charge of 271.9 ± 25.94 fC (mean ± SEM), which corresponds to ∼8.4 x 10^5^ molecules of 5-HT per vesicle (assuming two transferred electrons per oxidized molecule)^52^. We found that the spike charge generated from 5-HT released from single vesicles in EC cells is much larger than what has been reported for 5-HT release from small synaptic vesicles (2.9 fC charge; 45.6 nm diameter) or neuronal dense core vesicles (33.6 – 52.3 fC charge; 88.1 – 103 nm diameter)^49^. Our findings therefore indicate, that – assuming similar intravesicular 5-HT concentrations – EC cells store and release the vast majority of 5-HT from relatively large secretory vesicles.

Amperometric spike parameters such as rise time (0.642 ± 0.028 ms; mean ± SEM) and half-width (3.031 ± 0.147 ms; mean ± SEM) (Fig. 3g and h) provide information on the kinetics of the fusion event and the diffusion of the transmitter to the detection electrode. Similar to monoamine release from other secretory cells^53–55^, 5-HT release from EC cell vesicles was in many cases not an all-or-none event, but 83.84 ± 1.19% (mean ± SEM) of spikes per recording exhibited a so-called prespike foot signal (Fig. 3c, d, and i-l). This foot signal corresponds to the diffusion of the vesicular transmitter through a slowly opening fusion pore prior to full fusion of the vesicle with the plasma membrane^56^. Analysis of individual amperometric spikes also revealed a large heterogeneity especially with respect to the charge and amplitude of individual spikes (Fig. 3n-p).

We then employed our high-resolution secretion assay to test whether functional properties of 5-HT release differed between EC cells in acutely generated duodenal epithelial monolayer cultures (ACU) and EC cells in monolayer cultures derived from duodenal organoids (ORG) that have previously been in culture between 2 and 11 weeks (2-12 passages; Supplementary Fig. S2a-n). While organoids in culture can be propagated for long periods of time^57^, it is unknown how an extended period in culture may influence 5-HT metabolism and EC cell function. We performed this experiment by infusing cells with a solution containing a high (> 20 *µ*m) [Ca^2+^] to substantially trigger vesicle fusion, which came, however, at the expense of a higher number of overlapping amperometric spikes that were excluded from the analysis (see Methods). Comparing 5-HT secretion from EC cells in these two different preparations, we found only very subtle differences in the kinetics of 5-HT release (Supplementary Fig. S2).

Thus, our analysis revealed that EC cells in organoid cultures do not experience major cellular changes that would affect basic functional properties such as the amount of 5-HT stored and released from vesicles.

### The majority of 5-HT is released from EC cells with large dense-core vesicle (LDCV)-like slow kinetics

We then addressed the question whether 5-HT release from EC cells may occur with synapse-like, faster kinetics^49, 53^ than that of other monoaminergic systems that predominately employ LDCV-mediated exocytosis^58^. We therefore compared the kinetics of single vesicle 5-HT release events from EC cells to those of catecholamine (adrenaline and noradrenaline) induced amperometric spikes from functionally well-characterised adrenal chromaffin cells. To allow for a direct comparison, acute duodenal intestinal monolayer and isolated adrenal chromaffin cell cultures were generated from the same adult Tph1-CFP animal. We then analysed basic properties of amperometric currents recorded from EC cells (ECC; electrode potential +650 mV) and adrenal chromaffin cells (ACC; +800 mV) (Fig. 4a-o). Using this approach, we found only few differences in the amount of transmitter released from individual fusing vesicles such as that oxidation of secreted 5-HT from EC cells resulted in amperometric currents with a slightly larger average charge (Fig. 4d). The measured spikes further exhibited slower kinetics (Fig. 4f and g) than that generated by catecholamines from adrenal chromaffin cell.

**Figure 4.**
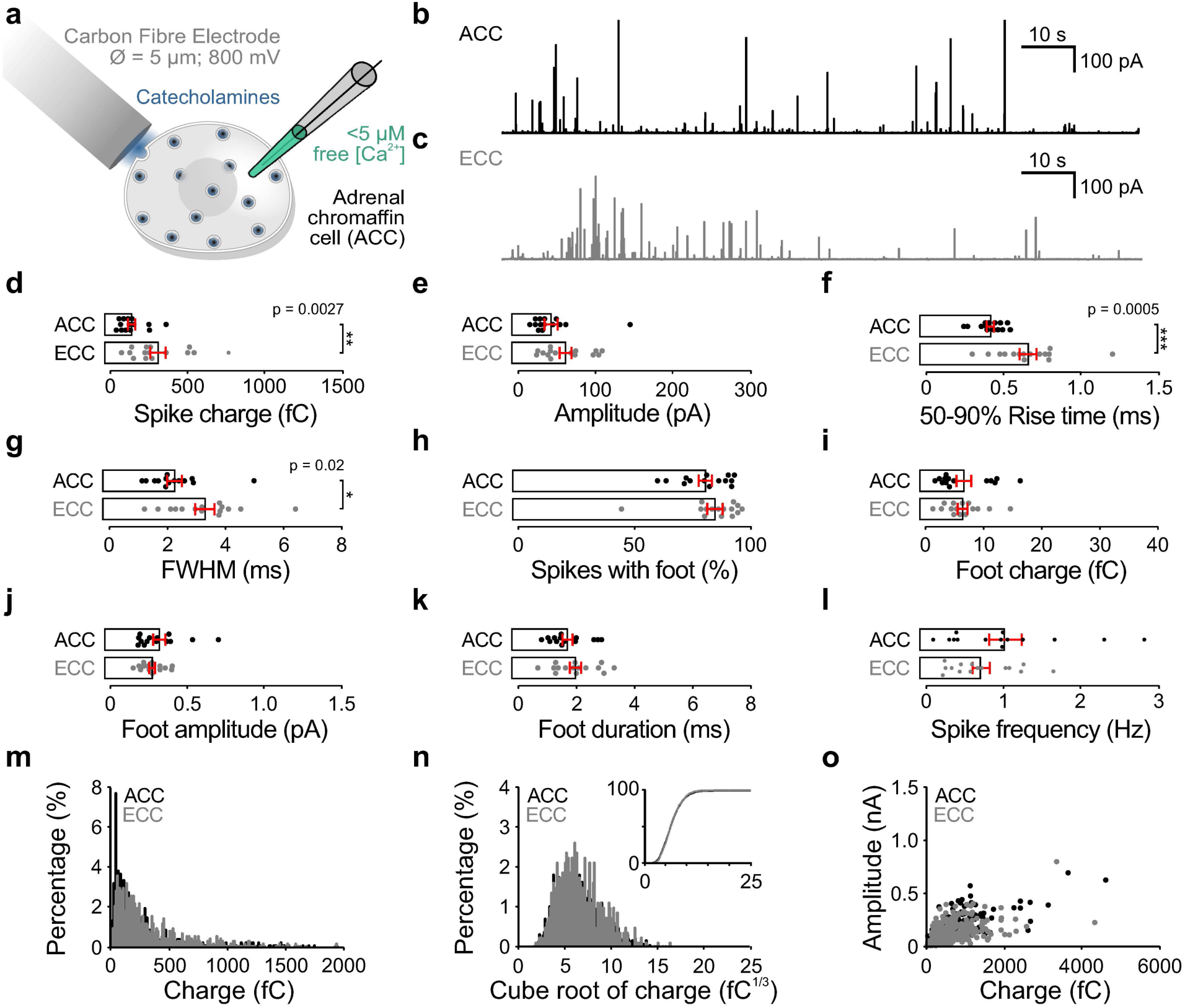
Comparison of Monoamine Release from Mouse EC Cells (ECC) and Adrenal Chromaffin Cells (ACC) by Carbon Fiber Amperometry. **a** Illustration of the experimental design. ACC and ECC cultures were generated from the same animals. Exocytosis in ACCs was triggered by infusing individual cultured cells with a solution containing moderate (< 5 *µ*m) [Ca^2+^]. Catecholamines released from single fusing vesicles were oxidized at the carbon-fiber electrode held at +800 mV. Serotonin release was measured as described in Fig. 3a. **b**-**c** Representative amperometric recording of ACCs and ECCs. **d** Median spike charge (fC). **e** Median maximum spike amplitude (pA). **f** Median 50-90% rise time (ms). **g** Median full-width half maximum (FWHM) corresponding to the width (ms) at half maximal amplitude. **h** Percentage of spikes with a foot signal from all spikes analyzed per cell (%). **i** Median foot charge (fC). **j** Median maximum foot amplitude (pA). **k** Median foot duration (ms). **l** Average spike frequency (Hz) per cell recorded during the 120 s recording interval before excluding overlapping spikes for the analysis of spike properties (**d**-**k**). **m** Frequency distribution of the amperometric spike charge (fC) of all events measured over all cells [bin width 10 fC, only events <2000 pA; ACC, 636 spikes, ECC, 783 spikes]. **n** Frequency distribution plotting the cube-root of amperometric spike charges (fC^1/3^) for all events measured over all cells [bin width 0.1 fC^1/3^; ACC, 645 spikes, ECC, 792 spikes]. Insert, cumulative frequency distribution for the same data. **o** Scatterplot illustrating the relationship between amperometric spike charge (fC) and maximum amplitude (nA) for each event measured [ACC, 645 spikes, ECC, 792 spikes]. Error bars indicate mean ± SEM; p < 0.05; **p < 0.01; ***p < 0.001. ACC, 2 cultures, 14 cells; ECC, 2 cultures, 15 cells.

Thus, our data indicate that the majority of 5-HT released from mouse EC cells is likely mediated by LDCVs that fuse with slow kinetics comparable to those of neuroendocrine cells.

### A high-throughput CLEM workflow for ultrastructural imaging of EC cells

To assess morphological features of vesicles present in EC cells, we next set up an experimental workflow permitting ultrastructural imaging of cultured EC cells. The fact that EC cells are found scattered at low density amongst other intestinal epithelial cell types, both *in vivo* as well as in the monolayer culture setting, necessitated a CLEM approach to rapidly find, as well as unequivocally identify EC cells, based on their genetic properties (Fig. 5).

**Figure 5.**
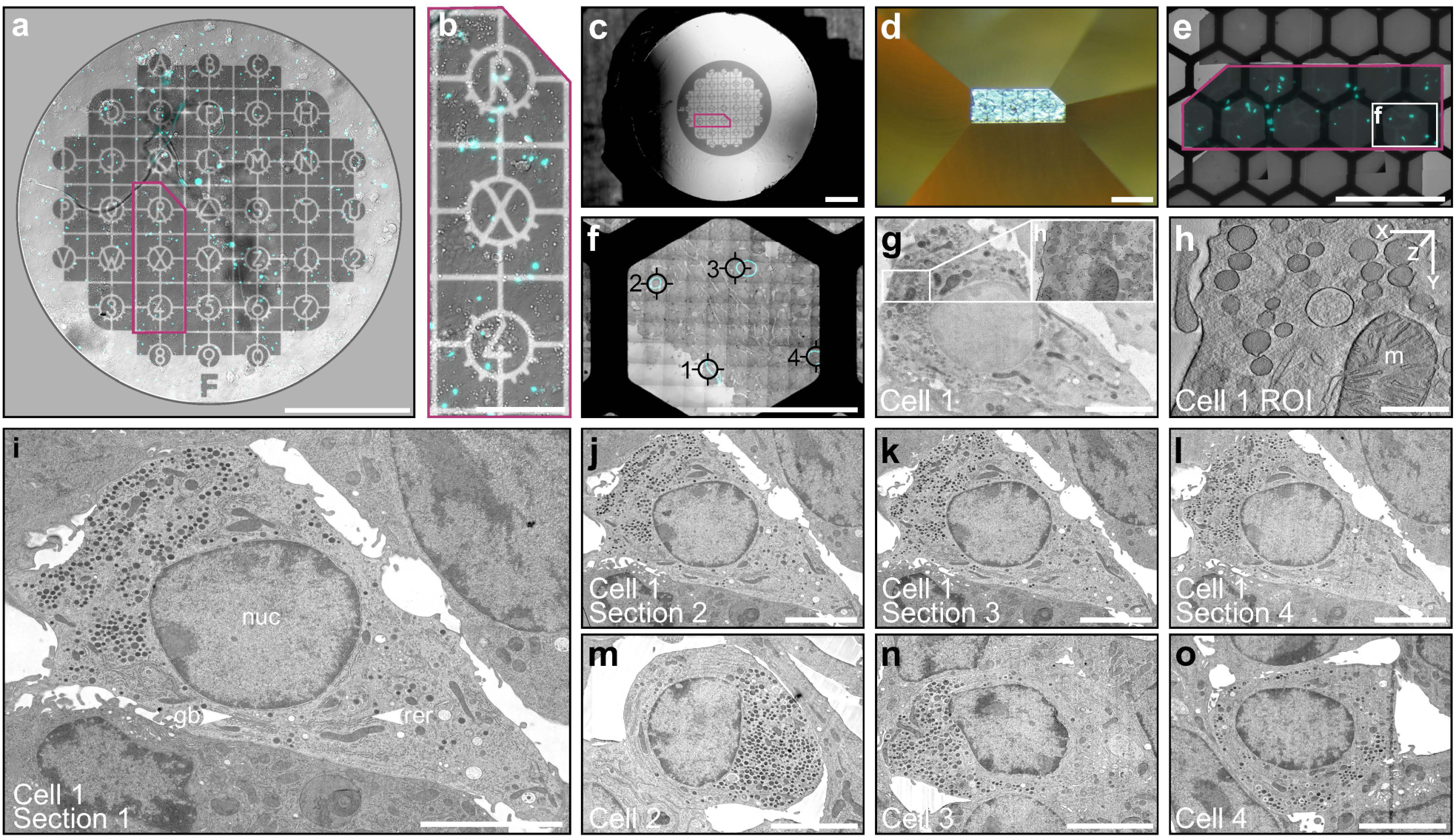
Workflow for Correlative Light and Electron Microscopy (CLEM) Experiments. **a** Epithelial monolayer cultures from Tph1-CFP mice were generated directly on sapphire discs decorated with a carbon coordinate system. On-sapphire cultures were subjected to live imaging to map the location of CFP-positive EC cells with respect to the coordinate system. **b** Coordinates of interest exhibiting high densities of CFP-positive EC cells were selected within an area (magenta outline) of dimensions compatible with transmission electron microscopy (TEM). **c**-**d** Plastic embedded culture viewed by reflected light before (**c**) and after (**d**) block-face trimming for targeted ultramicrotomy. Carbon coordinates remained on the surface of the epoxy resin block following the removal of the sapphire disc. **e**-**f** The light microscopic coordinates of CFP-positive cells superimposed upon plastic sections prepared for 2D and 3D ultrastructural analyses. Cyan outlines (**f**) indicate the predicted positions of CFP-positive cells visualized by live imaging and black ROIs mark the actual location of the cells shown in **g**-**o** (position 1: **g**-**l**; position 2: **m**; position 3: **n**; position 4: **o**). **g** 2D projection image of an EC cell in a 250 nm semithin section decorated with 10 nm gold fiducial particles. The insert indicates the region of interest selected for tilt-series acquisition and electron tomogram generation. **h** High-magnification electron tomographic subvolume showing secretory granules of different sizes (m, mitochondrion). **i**-**l** Serial ultra-thin section electron micrographs acquired through cross-section of the same EC cell used for electron tomography (**g** and **h**) (gb, golgi body; nuc, nucleus; rer, rough endoplasmic reticulum). **m**-**o** Electron micrographs acquired through identified CFP-positive cells from positions 2 (**m**), 3 (**n**), and 4 (**o**) similarly exhibit a polarized distribution of secretory granules. Scale bars: 1 mm (**a**, **c**); 250 *µ*m (**b**); 500 *µ*m (**d**, **e**); 50 *µ*m (**f**); 5 *µ*m (**g**, **i**-**o**); 500 nm (**h**).

To achieve this, we generated acute intestinal monolayer cultures from Tph1-CFP mice directly onto sapphire discs, which are compatible freezing substrates for various high-pressure freezing devices. To map the location of CFP-positive EC cells prior to fixation for electron microscopy, sapphire discs were decorated with a coordinate system made of a thin layer of carbon before starting the cultures (see Methods, Fig. 5a). We then performed live light microscopic imaging of 2D monolayer cultures to determine the position of individual cells expressing the fluorescent reporter with respect to the carbon coordinates. Based on the location of EC cells on the sapphire discs, regions exhibiting a high density of CFP-positive cells were identified (Fig. 5b). We then subjected the on-sapphire disc monolayer cultures to high-pressure freezing fixation followed by cryo-substitution and epoxy-resin embedding. This fixation method is known to be less prone to introduce artefacts that affect the cellular ultrastructure of neurosecretory cells, likely caused by classical chemical fixation approaches combined with room-temperature dehydration^59^. Sapphire discs were then removed from the plastic block surface, which exposed the clearly visible carbon coordinates (Fig. 5c) for guided trimming of the block surface and targeted ultramicrotomy (Fig. 5d). Using the light microscopic map of CFP-positive cells on the coordinate grid as a template (Fig. 5e) it was easily possible to locate the position of individual EC cells in sections imaged by transmission electron microscopy (TEM; Fig. 5f). We applied this workflow to map EC cells both in semithin sections intended for electron tomography, to accurately resolve sizes of single vesicles (Fig. 5g and 5h), and in ultrathin sections for TEM, to inspect cross-sections of entire EC cells (Fig. 5i-o). Although our CLEM workflow admittedly constitutes a rather crude approach of mapping cells for ultrastructural analyses (i.e. no additional fiducial markers were applied that would increase the spatial accuracy of the method), we experienced a very high success rate in finding EC cells for ultrastructural imaging at light microscopically predicted sites. The reason for the high reliability of the approach can be attributed to: i) EC cells and other EEC types comprise the absolute minority of all intestinal epithelial cells and are therefore found spatially segregated (e.g. Fig. 5b), and ii) EECs can clearly be distinguished from other surrounding cell types in culture such as enterocytes based on morphological criteria, such as the presence of LDCVs (Supplementary Fig. S3a-d). We rarely found other EECs exhibiting LDCVs in very close proximity to the CFP-positive cells.

### EC cells harbour only few small clear-core vesicles

We then approached the question of whether EC cells indeed store 5-HT predominantly in LDCVs as indicated by the functional single-cell electrochemistry experiments (Fig. 3, Fig. 4**, and** Supplementary Fig. S2). Firstly, we found by inspecting electron micrographs of ultrathin EC cell cross-sections, that EC cells in monolayer epithelial cultures retained their polarity and concentrated the vast majority of secretory vesicles at one side of the cell (Fig. 5i-o). Secondly, to obtain a precise readout of vesicle sizes, we performed electron tomography on semithin sections of cytoplasmic EC cell regions that contained a high density of vesicles (Fig. 6a). The main advantage of this imaging approach in comparison to analyses of vesicle sizes in 2D projection images is that the diameter of individual vesicles can be measured directly at its largest extent in three-dimensional tomographic reconstructions (Fig. 6a-c). This is particularly advantageous for the accurate assessment of LDCV dimensions in neuroendocrine cells that often exhibit a large variability in vesicle diameters that can range between several hundred nanometers^51, 60, 61^. Classical analysis of electron micrographs taken of ultrathin sections (∼40-60 nm thickness) would therefore result in a systematic underestimation of actual vesicle diameters unless a correction for section thickness is applied^49, 60^.

**Figure 6.**
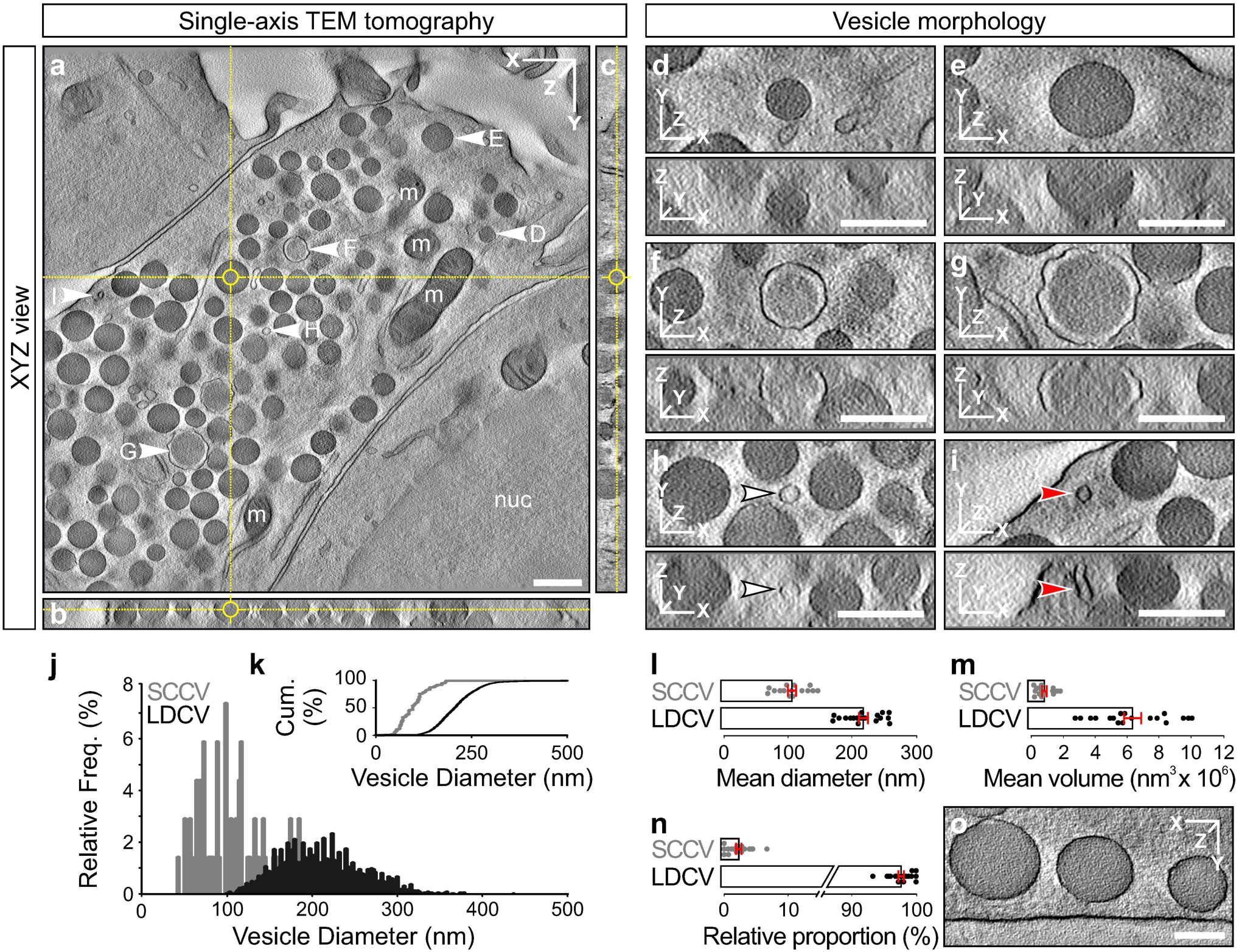
Ultrastructural Analysis of Secretory Vesicles in Mouse EC Cells by Single-Axis Electron Tomography. **a**-**c** High-magnification electron tomographic subvolume of an EC cell enabling accurate measurement of vesicle dimensions in 3D at the widest diameter (“midline”) of the vesicle. Individual vesicles exhibited a heterogeneity in size and appearance (**d**-**h**) (nuc, nucleus; m, mitochondria). **d**-**g** Vesicles were scored as large dense-core vesicles (LDCVs) based on the presence of an electron dense vesicle core. **h** Vesicles were scored as small clear-core vesicles (SCCVs; white arrowhead) when no apparent intravesicular structures were visible. **i** Tubular structures (red arrowhead) that appear spherical in 2D projections could be identified and excluded from the analysis. **j** Frequency distribution of vesicle diameters separated by SCCVs and LDCVs with respect to the total number of vesicles analyzed for the respective groups [bin width 2 nm; LDCVs, 2724 vesicles, SCCVs, 67 vesicles]. Data pooled over all tomograms analyzed. **k** Cumulative frequency distribution of the data shown in (**j**). **l** Mean vesicle diameter (nm). **m** Mean vesicle volume (nm^3^ x 10^6^). **n** Relative proportion of SCCVs and LDCVs per tomogram (%). **o** High-magnification electron tomographic subvolume showing three LDCVs in proximity to, but not in direct contact with, the plasma membrane. Scale bars: 500 nm (**a**); 200 nm (**d**-**I**, **o**). Error bars indicate mean ± SEM. 2 cultures, 18 tomograms, 2791 vesicles.

Similar to well-characterised adrenal chromaffin cells^51^, we found that EC cell vesicles exhibited a large heterogeneity in size and in the compactness of the dense-core (Fig. 6d-6h). For subsequent analyses, we separated vesicles into two groups based on their core appearance, namely LDCVs (Fig. 6d-6g) and small clear-core vesicles (SCCVs; Fig. 6h). Vesicular structures were assessed in three-dimensional tomographic volumes in both, the x-y and x-z dimension, to exclude structures from the analysis that were indeed tubular in nature and therefore likely did not constitute secretory organelles (Fig. 6i). Assessing the relative size distributions for LDCVs and SCCVs confirmed that SCCVs are on average smaller [diameter (∅): 105.9 ± 6.303 nm; mean ± SEM) than LDCVs (∅: 216.9 ± 6.901 nm; mean ± SEM; Fig. 6j-l **and** Supplementary Fig. S3e], which corresponds to a 7.68-fold difference in the average vesicle volume (Fig. 6m). When analyzing the relative contribution of each vesicle type per tomogram (Fig. 6n), we found that SCCVs only constituted on average 2.3% of all vesicles analysed. Of note, adrenal chromaffin cells exhibit a large number of secretory granules in direct contact with the plasma membrane in a membrane-attached or “docked” state^51^. This morphological vesicle state corresponds in neuronal synapses to a pool of highly fusogenic vesicles, which are primed for rapid merger with the plasma membrane in response to stimulation and are major determinants of the speed of synaptic transmission^62, 63^. In EC cells, however, we observed that only extremely few granules (1 SCCVs and 5 LDCVs in 18 tomograms) were found in direct contact with the plasma membrane (Fig. 6o).

In summary, our ultrastructural data indicate that the majority of 5-HT in EC cells is likely stored in LDCVs and not SCCVs. This finding in line with our data showing that the kinetics and amount of 5-HT released from single vesicles as assessed by single-cell carbon fibre amperometry, resemble those in other endocrine systems (Fig. 4).

### Key components of the molecular transmitter release machinery expressed in EC cells

In secretory cells, Ca^2+^-triggered fusion of peptide hormone or neurotransmitter storing vesicles with the plasma membrane is mediated by a specialized and highly conserved molecular machinery. The cell-specific composition and subcellular organisation of this machinery are important determinants of the efficacy of secretion. For instance, presynaptic regulators of neurotransmitter release in neuronal synapses i) control the speed of information transfer at synaptic cell-to-cell contact sites through the presence of a pool of highly fusogenic or “primed” vesicles^64^, ii) delineate dedicated plasma membrane regions or “active zones” to spatially restrict the fusion of vesicles and the concomitant release of signalling molecules^65, 66^, iii) allow rapid and purposeful adaptation of the efficacy of transmitter release during phases of neuronal activity^67^, and iv) regulate the temporal precision of excitation and secretion coupling and the sensitivity of the vesicle fusion reaction to changes in the intracellular [Ca^2+^]^68^. Determining the molecular mechanisms by which Ca^2+^-mediated release of serotonin from EC cells is executed, will therefore help to identify molecular targets for pharmacological interventions to fine-tune serotonin signalling in the gut for the treatment of EC-cell associated diseases.

To resolve EC-cell specific expression of key known molecular mediators of exocytosis in neuronal synapses, we first performed a gene expression analysis of pooled fluorescence-activated cell sorting (FACS) purified small intestinal EC cells^36^ (Fig. 7a-i). The RNA samples used here were generated from enzymatically digested mouse gut tissue biopsies, from which EC cells were isolated by FACS with the help of a primary antibody directed against 5-HT and a secondary antibody coupled to the fluorophore Alexa488^36^. As described previously, sorting of cells at 488nm-excitation resulted in a very pure EC cell population as determined by the expression of the EEC marker Chromogranin A (ChgA) and the EC cell marker Tph1 as compared to the non-excitable cell population^36^ (Fig. 7a-c). We performed RT-PCR on isolated RNA samples using a custom-designed qPCR array for various components of the presynaptic neurotransmitter release machinery (Fig. 7d-i). Employing this approach, we could identify candidate genes transcripts that were highly expressed in the FACS-purified EC cell population. From this list, we picked several candidates for subsequent analyses: SNAREs: SNAP-25, and VAMP2 (Fig. 7d); Vesicle priming proteins: CAPS-1 (Cadps) and Munc13-1 (Unc13a; Fig. 7e); and Monomeric G-proteins: Rab3s (Fig. 7g).

**Figure 7.**
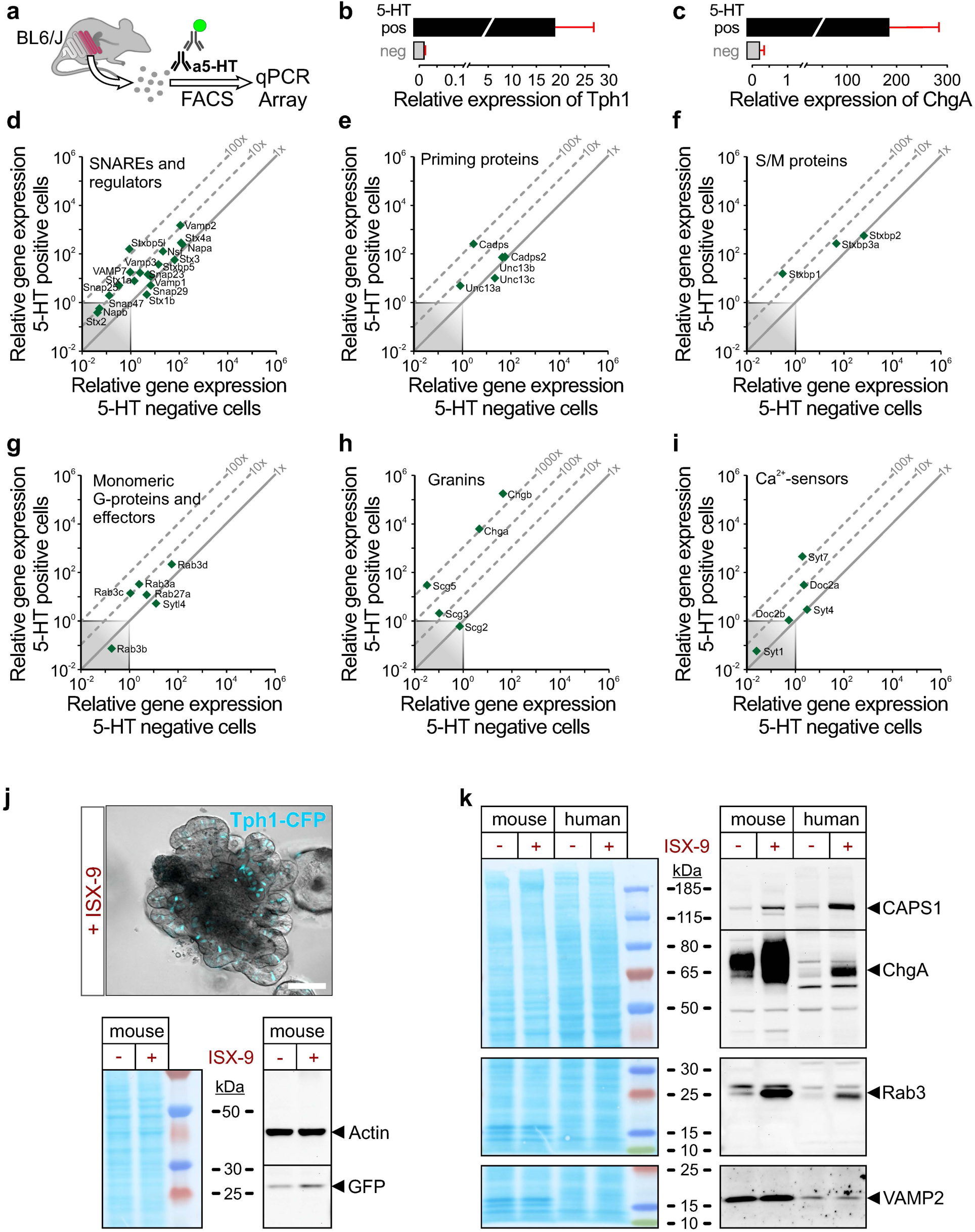
Characterization of the Expression of Candidate Components of the Presynaptic Neurotransmitter Release Machinery in EC cells by RT-PCR and Protein Immunoblotting. **a** 5-HT antibody-mediated fluorescence-activated cell sorting (FACS) purification of EC cells from the small intestine of wildtype mice, RNA extraction and qPCR. The qPCR analyses presented here were performed on RNA isolated during a previous in-house study^1^. **b**, **c** qPCR analyses of the expression of the EC marker genes tryptophan hydroxylase 1 (Tph1)(**b**) and Chromogranin A (ChgA)(**c**) in 5-HT positive and 5-HT negative fractions of sorted cells (the same data were shown in Fig. 1 of^36^). **d**-**i** qPCR analysis expression data for known components of the presynaptic molecular neurotransmitter release machinery divided into sub-categories (**d**, SNAREs and regulators; **e**, priming proteins; **f**, S/M proteins; **g**, Monomeric G-proteins and effectors; **h**, granins; **i**, Ca^2+^-sensors). The 45°-angled dotted lines display the fold change enrichment in 5-HT positive cells compared to 5-HT negative cells. The grey-shaded square marks what was considered as noise. **j** Organoid generated from a Tph1-CFP mouse and subjected to the 48 h ISX-9 pulse protocol^69^ (top panel). Western blot analysis of duodenal organoids from Tph1-CFP mice treated with ISX-9 (+) or DMSO only (-) using antibodies directed against Actin (loading control) and GFP to detect CFP (Lower panel). **k** Western blot analysis of duodenal organoids from Tph1-CFP mice and human ileal organoids treated with ISX-9 (+) or DMSO (-) using antibodies directed against CAPS-1, Chromogranin A (ChgA), Rab3 proteins, and VAMP2. Protein stain of the nitrocellulose membrane (MEM Code, left panel) and chemiluminescent signals detected for the same membrane segments after immunolabelling with different antibodies (right panels). Scale bar: 100 *µ*m (**j**).

### CAPS1 and Rab3 Presynaptic Proteins are Enriched in Mouse and Human EC cells

To confirm whether we could detect the candidates identified from the qPCR screen also on the protein level in EC cells, we used a combination of Western blot analysis of EC-cell enriched organoid cultures (Fig. 7j-k**, and** Supplementary Fig. S4), as well as immunocytochemistry (Fig. 8a-d, **and** k-n) and immunohistochemistry (Fig. 8e-j, **and** o-t) for light microscopic analysis of EC cells in 2D monolayer cultures and gut tissue, respectively.

**Figure 8.**
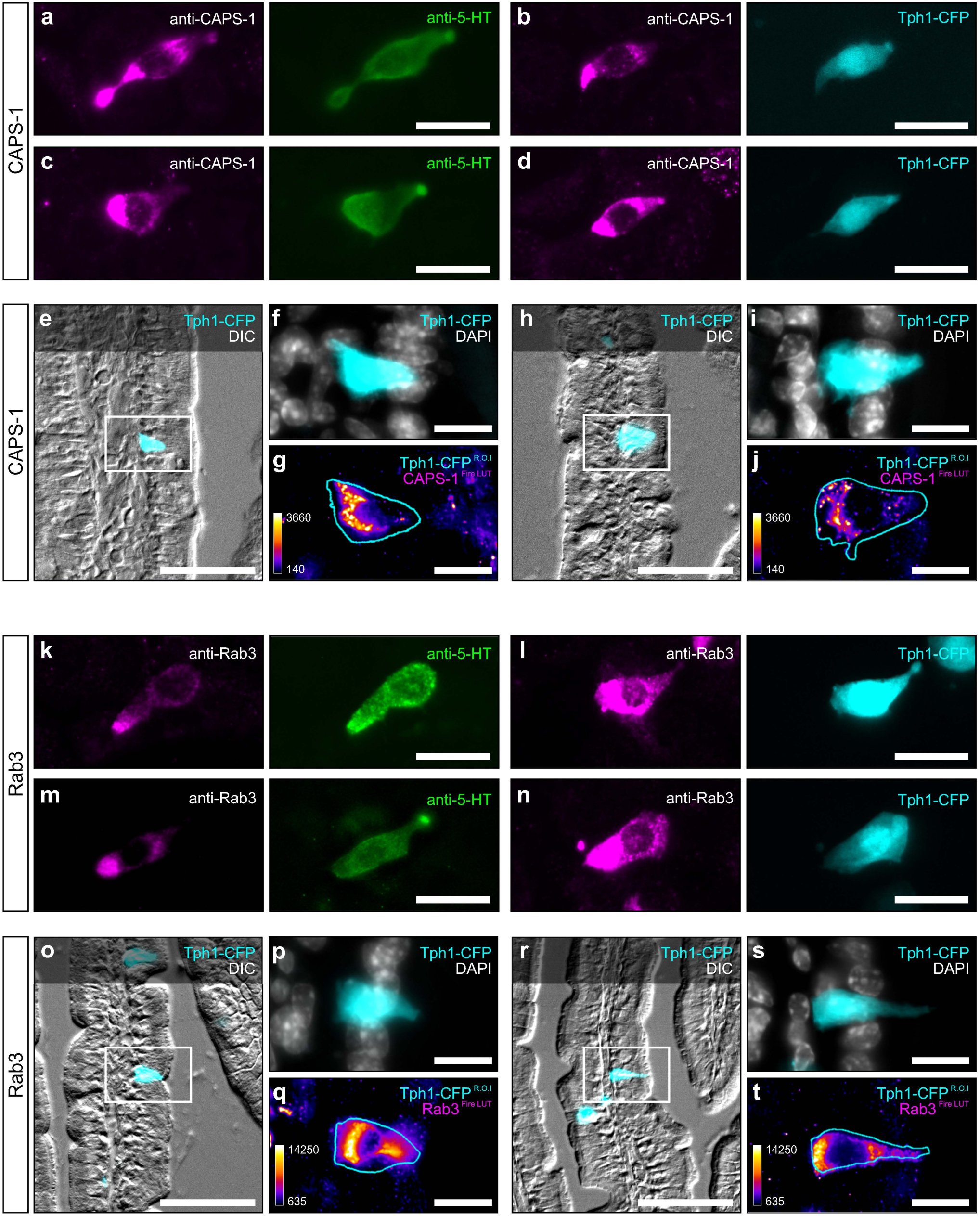
Immunofluorescent Labelling of Key Molecular Components of the Presynaptic Neurotransmitter Release Machinery in EC cells. **a**-**d** EC cells in intestinal epithelial monolayer cultures of Tph1-CFP mice exhibiting endogenous CFP-fluorescence (cyan) and were stained with antibodies against 5-HT (green) and CAPS-1 (magenta). **e**-**j** EC cells in tissue of the proximal small intestine of Tph1-CFP mice exhibiting endogenous CFP-fluorescence (cyan, **e, f**, **h**, **i**; cyan outline in **g** and **j**) were stained with an antibody directed against CAPS-1 (fireLUT, **g** and **j**) and DAPI (white, **f** and **i**). **k**-**n** EC cells in intestinal epithelial monolayer cultures of Tph1-CFP mice exhibiting endogenous CFP-fluorescence (cyan) were stained with antibodies against 5-HT (green) and Rab3 proteins (magenta). **o**-**t** EC cells in tissue of the proximal small intestine of Tph1-CFP mice exhibiting endogenous CFP-fluorescence (cyan, **o, p**, **r**, **s**; cyan outline in **q** and **t**) were stained with an antibody against Rab3 proteins (fireLUT, **q** and **t**) and DAPI (white, **p** and **s**). Panels **a-d** and **k**-**n** show z-projections (sum slices) of an image stack, and panels **e**-**j** and **o**-**t** show single image planes acquired from a z-stack. Scale bars: 20 *µ*m (**a**-**d** and **k**-**n**), 50 *µ*m (**e**, **h**, **o**, and **r**), and 10 *µ*m (**f**, **g**, **i**, **j**, **p**, **q**, **s**, and **t**).

To increase the proportion of EC cells in mouse organoids for biochemical analyses (Fig. 7j-k**, and** Supplementary Fig. S4), we treated small intestinal organoid cultures derived from Tph1-CFP mice with the small molecule isoxazole-9 (ISX-9)^69^. Exposure to ISX-9 activates a transcriptional program involving neurogenic differentiation 1 (NeuroD1) that promotes pancreatic β-cell differentiation and insulin production^70^ as well as EEC differentiation with a specific bias towards 5-HT positive EC cells^69^. In our experimental system, increased CFP protein levels were detected in culture homogenates of EC-cell enriched (ISX-9 treated) organoids compared to the control (DMSO vehicle treated) sample as determined by the use of an antibody directed against GFP for immunoblotting (Fig. 7j). Our results reproduce previous findings showing that the proportion of EC cells in mouse and human intestinal organoids is increased by the ISX-9 stimulation protocol^69^.

We next assessed whether candidate proteins of the presynaptic release machinery were specifically enriched in homogenates of ISX-9 treated mouse and human^69^ organoids. Using primary antibodies specific for CAPS-1 (SySy, Cat# 262 003) and all Rab3 isoforms (a, b, c, d; SySy #107 003), our approach revealed a prominent enrichment of CAPS-1 and members of the Rab3 protein family together with the EEC marker ChgA in homogenates from ISX-9 stimulated mouse and human organoids (Fig. 7k). Interestingly, while we were able to detect the vesicular SNARE protein VAMP2 in mouse and human organoid homogenates, no specific enrichment was observed as a result of ISX-9 treatment (Fig. 7k). These data provide strong evidence that CAPS-1 and Rab3 proteins are expressed in EC cells, while VAMP2 expression in the gut epithelium is likely not specific for EC cells or even EEC. Indeed, it has previously been shown that VAMP2 mRNA is expressed together with VAMP8 in mucin secreting goblet cells of mouse colonic organoid cultures^71^. Using immunoblotting, we failed to detect immunoreactive protein bands at the expected sizes for any of the sampled conditions using antibodies directed against SNAP-25 (Supplementary Fig. S4a) and Munc13-1 (Supplementary Fig. S4b), indicating that these proteins are either not at all expressed in gut organoid samples, or at levels below the detection threshold. In line with this finding, SNAP-25 protein was previously also not detected in colonic organoid homogenates^71^. Future studies will have to address the discrepancy between the failed detection of SNAP-25 and Munc13-1 in organoid homogenates by Western Blot analyses (Supplementary Fig. 4) despite the prominent mRNA expression levels in FACS purified EC cells (Fig. 7d and e; see also^17^).

Next, we set out to assess the subcellular localization of our proteins of interest in mouse small intestinal EC cells. Understanding the organization of the molecular release machinery (i.e. presence of synapse-like presynaptic specializations or “active zones”) is important to answer the question whether EC cells form synapse-like connections with neuronal processes. Using isoform-specific antibodies for light microscopic analysis, we found that CAPS-1 and Rab3 immunoreactivity was readily detected in Tph1-CFP-positive EC cells in 2D monolayer cultures (Fig. 8a-d **and** k-n) and in gut tissue sections (Fig. 8e-j **and** o-t). The immunofluorescent signal for CAPS-1 (Fig. 8a-j) and Rab3 proteins (Fig. 8k-t) was found to be concentrated in the cytoplasm at the basolateral side, indicative of an association of these proteins with secretory vesicles.

In conclusion, we demonstrate that key regulators of the neuronal molecular transmitter release machinery are expressed on the protein level in EC cells. Based on their known function in synapses^72–75^ and other secretory systems^50, 76, 77^, it is tempting to presume a prominent role for these factors in mediating 5-HT secretion in the gut.

## Discussion

We present here an experimental approach to monitor Ca^2+^-triggered 5-HT release from genetically identified cultured EC cells at single-vesicle resolution *in vitro*. To achieve this, we performed carbon-fibre amperometry for the electrochemical detection of 5-HT after infusion of EC cells with moderate and high [Ca^2+^] containing solutions. In contrast to previous results obtained from amperometric recordings performed on isolated EC cells showing that the kinetics of 5-HT release are extremely fast (∼0.1 ms half-width)^39^, we demonstrate here that the majority of 5-HT is released with relatively slow kinetics (∼3 ms half-width; Fig. 3h) that are comparable to those known for other neurosecretory systems that employ LDCVs (Fig. 4).

A variety of different preparations, stimulation protocols, and 5-HT detection assays have been employed in the past to monitor monoamine secretion from EC cells in different physiological contexts:

i. Enzyme-linked immunosorbent assay (ELISA) to measure 5-HT secreted into solution from cultured EC cell-like cancer cells including BON-cells^78, 79^, primary monolayer cultures^23^, 3D intestinal organoids^69^, enzymatically digested and density gradient-centrifugation purified EC cells^39, 80–82^, as well as tissue biopsies^5, 79, 82, 83^,
ii. high-performance liquid chromatography (HPLC) applied to the supernatant of stimulated gut tissue biopsies^36^,
iii. 5-HT_3_ receptor-expressing HEK “biosensor” cells to detect 5-HT released from nearby EC cells in organoids^17^ and monolayer cultures^6^ using whole cell patch clamp electrophysiology or Ca^2+^-imaging, and
iv. carbon fibre amperometry for the detection of bulk 5-HT release from gut tissue samples^36, 84–86^, and single-cell carbon fibre amperometry for the study of 5-HT secretion from enzymatically digested and density gradient-centrifugation purified EC cells^39, 40, 84^ and BON cells^87^.

All of these 5-HT detection assays differ substantially with respect to their sensitivity, spatial precision, temporal resolution, and application. For instance, analyses by ELISA or HPLC provide no spatial and little temporal information with respect to release from individual cells as samples are generally stimulated for a period of time to evoke sufficient 5-HT release into the extracellular medium. Carbon fibre amperometry applied to tissue patches enables the direct detection of 5-HT released from many EC cells simultaneously, and the measured amperometric currents therefore correspond to bulk secretion of 5-HT. This approach offers high sensitivity and temporal resolution, but it cannot provide precise spatial information. However, none of these three techniques rely on genetic approaches to label individual EC cells and they therefore have the advantage to be applicable to many different experimental systems including human gut tissue biopsies. In contrast, biosensor experiments offer a high spatial and temporal resolution for the detection of 5-HT, which is largely determined by the diffusion of the transmitter from the EC cell to the biosensor cell. It is, however, important to note that the measurement of 5-HT3 receptor currents in HEK cells can only be interpreted as an indirect approximation for the total amount of 5-HT released from a given cell. This is due to the fact that these measurements depend on the receptor expression levels in the biosensor cell. In comparison, our single-cell carbon fibre amperometry approach provides an extremely high temporal resolution to monitor 5-HT secretion from EC cells in near real time. Moreover, the direct detection of 5-HT permits a highly standardized analysis of the amount of 5-HT released from individual vesicles and of the kinetics of this process. Finally, we anticipate that our experimental system will be ideally compatible with recently developed functional 5-HT imaging approaches using genetically encoded 5-HT-sensors^88^ or exogenously applied nanosensors^89^, which would allow to resolve potential “hotspots” for 5-HT release in EC cells. To enable EC cell-specific expression of genetically encoded sensors as well as opto-or chemogenetic actuators in culture, we have generated a novel inducible Tph1-iCreERT2 mouse line (Supplementary Fig. S5). That said, one caveat of our method in general is that it indeed requires the help of genetic strategies such as the use of the Tph1-CFP transgenic mouse line to discriminate EC cells from other epithelial cells in culture. However, recent advantages in combining novel gene editing technologies with human intestinal organoid cultures^90, 91^ that can then be converted to monolayer cultures for functional assays, will make it possible to study 5-HT release from EC cells in different physiological and disease contexts. We therefore envision our experimental approach as an ideal system to probe the cell-specific effects of pharmacological approaches designed to modulate 5-HT metabolism in EC cells, such as novel molecular Tph1-inhibitors^92, 93^.

Using single-cell carbon fibre amperometry, we observed that EC cells exhibited a large percentage of amperometric spikes with a prominent foot signal. This finding will have implications for our understanding of how 5-HT and peptide hormones co-expressed in EC cells might be released from the same vesicle. Foot signals are caused by diffusion of transmitters through membrane fusion-pores during the early stages of the membrane fusion reaction. Vesicular fusion-pores are dynamic structures that can either prelude i) full-collapse vesicle fusion and degranulation, or ii) fast membrane recycling events during which the fusion pore only transiently opens before proceeding to so-called kiss-and-run endocytosis^94^. Several studies have proposed that during mild stimulation, transient kiss-and-run vesicle fusion may serve as the predominant secretion mode in adrenal chromaffin cells, whereas strong stimulation paradigms induce a shift towards full-collapse fusion^95, 96^. Large (neuro)peptides and hormones, which are often co-stored with smaller signalling molecules such as ATP^14, 97, 98^, serotonin^99^, and catecholamines^100^ in the same secretory vesicle, cannot diffuse through small initial fusion pores < 4nm^101^ and therefore require the presence of large, stable pores or full-collapse fusion. Vesicular fusion pores are therefore important structures that permit fine-tuning of the amount and the type of signalling substance released from a given vesicle. Conditions that cause defects in fusion-pore expansion in pancreatic β-cells have been shown to affect glucose-induced insulin secretion from these cells, which can be directly linked to the pathogenesis of type 2 diabetes^97, 102–104^. Understanding the kinetics of single vesicle fusion events in EC cells may therefore likewise contribute to a deeper understanding of EC cell-associated disease pathologies. Moreover, it would be interesting to test in the future whether 5-HT release kinetics change throughout EC cell maturation, a process which is marked by differential co-expression of the peptide hormones substance-P and secretin^30, 31, 33^. An important consideration is that our study applied a “brute-force” approach to trigger regulated exocytosis by infusing EC cells with solutions containing relatively high [Ca^2+^]. However, EC cells express voltage-gated sodium channels that render them electrically excitable^17, 23^, and they are also known to respond to various stimuli including nutrients, metabolites, irritants, and mechanical stimulation through intracellular signalling cascades and second messengers^17, 36, 80, 82, 105^. At least some of these stimulants trigger 5-HT secretion likely independent of action-potential firing^17^. In the future, it will therefore be important to test how the kinetics of 5-HT release change in response to different stimulation paradigms and secretagogues.

Using carbon-fibre amperometry, we found in our experimental system that EC cells release the majority of 5-HT from LDCVs rather than smaller clear-core vesicles. It is important to note, however, that we cannot rule out the possibility that we underestimate the contribution of 5-HT released from small synaptic-like vesicles due to experimental constraints. Firstly, LDCVs in adrenal chromaffin cells in different preparations including dissociated cultures are found all along the plasma membrane, whereas cultured EC cells retain their polarity and concentrate the majority of LDCVs at the basal side (Fig. 5i). Therefore, the placement of the carbon fibre during amperometric recordings has a huge impact on parameters such as the spike frequency. Secondly, for the analysis of amperometry data, we used a spike detection threshold of 5 pA^48^. Using this approach, we therefore likely miss monoamine release events that would have resulted from the fusion of small synaptic-like vesicles resulting in much smaller current spikes with faster kinetics^49, 53^. However, our functional data correlated well with our ultrastructural findings demonstrating that in genetically identified EC cells, the majority of vesicles are LDCVs with an average diameter of 217 nm (Fig. 6). The here reported sizes of LDCVs in cultured mouse EC cells are similar to those storing catecholamines in newborn mouse adrenal chromaffin cells^51, 106^. This is in line with our observation that basic amperometric spike properties are largely comparable between these two secretory cell types (Fig. 4). That said, previous work performed on acutely isolated guinea pig and human EC cells indicated that 5-HT is released with much faster kinetics than what we reported here for mouse EC cells from monolayer cultures^39^. Future studies will have to determine what causes these differences in the kinetics of 5-HT release between preparations.

Combining high-pressure freezing fixation, automated freeze substitution, and electron tomography on plastic sections allowed us to accurately assess vesicle morphologies and sizes in individual and genetically identified cultured EC cells using a CLEM approach. We found that the majority of vesicles in EC cells are LDCVs rather than SCCVs in line with our functional data, and that very few vesicles are found in close contact or “docked” to the plasma membrane. In the past, pre-and postembedding immuno-EM using antibodies directed against 5-HT revealed that 90% of 5-HT-immunoreactivity localizes to LDCVs in EC cells, whereas only ∼10% is present in the EC cell cytoplasm^107–109^, supporting the notion that the majority of 5-HT is stored in LDCVs in these cells. One potential caveat of our high-resolution imaging approach is that we focused only on small subcellular regions in individual EC cells and did not reconstruct entire cells. It is possible that EC cells form dedicated transmitter release sites that cluster small synaptic-like vesicles reminiscent of so-called “active-zones” at neuronal synapses in the brain, which we may have missed. Likewise, we have to consider the possibility that such putative release sites may only form in culture in the presence of a “postsynaptic” neuronal partner, which is absent in the 2D monolayer culture system. In line with previous findings^17^, we confirm here that EC cells express a large number of components that constitute the neurotransmitter release machinery at neuronal synapses in the brain. These data provide support for the notion that EC cells employ a similar molecular machinery as neurons to mediate secretory vesicle fusion. Moreover, several studies have reported the presence of neuron-like processes in EC cells *in vivo*^34, 110–113^, however, ultrastructural evidence for the presence of bona fide synaptic contact points between EC cells and neuronal processes is still lacking^13, 114^.

Finally, we have focused our functional and ultrastructural analyses in the present study on EC cells of the proximal small intestine. In this gut region, EC cells are involved in mediating *in vivo* nutrient-triggered rapid signalling towards the brain^24^, likely via activation of vagal sensory afferents that express the ionotropic 5-HT_3_ receptor^115^ and are found anatomically in close proximity to epithelial EC cells^17^. However, EC cells are distributed throughout the entire gastrointestinal tract^4^. 5-HT released from colonic EC cells modulates gut motility and serves as a pro-inflammatory factor. The composition of the gut microbiome of the large intestine has been shown to impact colonic 5-HT levels and thereby EC function, likely by modulating 5-HT metabolism in EC cells^83^. EC cells of the small intestine and colon also exhibit differences in their structural features^34, 35^ and in the molecular receptor repertoire with respect to their localization in the GI tract^36–38^. In the future, it will therefore become important to investigate whether the amount and kinetics of 5-HT release, EC cell ultrastructure, and molecular mediators of the 5-HT secretion process differ based on their location within the GI tract.

## Acknowledgements

This project has received funding from the European Union’s Horizon 2020 research and innovation programme under the Marie Sklodowska-Curie grant agreement No 843494 (C.I.) and the work was supported with funding provided by the Schering Stiftung („Boost your research“; C.I.), the Lundbeck Foundation (R307-2018-2973; C.I.), and the MSc/PhD program “Molecular Biology” – International Max Planck Research School at the Georg August University Göttingen (F.M.). B.H.C and C.I. hold a grant from the Schram-Stiftung. We are extremely grateful to Nils Brose for his continuous, generous, and unconditional support. We thank H. Taschenberger and C. Thomas for sharing analysis routines, M. Schwark and the DNA Core Facility for excellent technical support, P. Fonesca Pedro and A. Tsakmaki for the human organoids, H. Clevers for sharing the L-Wnt3A cell line, A. Lumsden, A. Martin, D. Goldspink, and V. Lu for sharing their expertise in culture techniques, U. Fünfschilling and M. Schindler of the MPIEM Transgene Facility for mouse embryology and zygote injections, J.B. Sørensen for helpful comments on the manuscript, and the members of the MPI-EM animal facility for mouse husbandry.

## Author Contributions

Conceptualization, C.I.; Methodology, A.S., B.H.C., and C.I.; Formal Analysis, A.S., B.H.C., F.M., V.S., M.L.L., and C.I.; Investigation, A.S., B.H.C., F.M., M.L.L., S.B., C.H., M.C., F.B., and C.I.; Resources, V.S., G.A.B., and F.B.; Writing – Original Draft, C.I.; Writing – Review & Editing, all authors; Visualization, B.H.C., F.M., M.L., and C.I.; Supervision, B.H.C., G.A.B., D.J.K., F.B., and C.I.; Funding Acquisition, C.I..

## Methods

### Resource Availability

#### Lead Contact

Further information and requests for resources and reagents should be directed to and will be fulfilled by the lead contact Dr. Cordelia Imig.

### Materials Availability

The newly generated Tph1-P2A-iCreERT2 mouse line will be made available by the authors upon publication and reasonable request.

### Data and Code Availability

Any computer-generated codes and raw data supporting the current study is available from the Lead Contact upon reasonable request.

## Experimental Model and Subject Details

### Animals

Mouse breeding and all experimental procedures involving primary cell culture experiments were carried out with permission of the Niedersächsisches Landesamt für Verbraucherschutz und Lebensmittelsicherheit (LAVES). Transcardial perfusion of mice was performed under the LAVES license 33.19-42502-04-18/2756. Animals were kept according to the European Union Directive 63/2010/EU and ETS 123. Mice were housed in individually ventilated cages (type II superlong, 435-cm2 floor area; TECHNIPLAST) under a 12 h light/dark cycle at 21 ± 1 °C with food and water ad libitum. The health status of the animals was checked regularly by animal care technicians and a veterinarian. The generation of the Tph1-iCreERT2 mouse line was performed under the LAVES license 33.19-42502-04-16/2100 as described below and crossed with C57BL/6 and Ai32 knockin (KI) mice^116^ to generate mice heterozygous for both, the Tph1-iCreERT2 and the Ai32 KI alleles. The 2D monolayer culture was generated from a 7-week-old male mouse. The Tph1-CFP line^29^ was imported from The Jackson Laboratory (Stock No. 028366) on a mixed genetic background (B6SJLF1/J and CD1). The line was rederived at the Max-Planck Institute of Experimental Medicine (Göttingen, Germany) and backcrossed for several generations with C57BL/6 mice. To compare monoamine release from adrenal chromaffin cells and EC cells measurements from both cell types were performed on cultures generated from the same 17-32 weeks old adult Tph1-CFP transgenic animals. Small intestine 3D organoid cultures were generated from 15-32 weeks old adult Tph1-CFP mice. For all experimental groups, mice of both genders were used, and the data were pooled. We did not check for significant changes between these conditions but report this information in **Supplementary Table S1**. For immunocytochemistry and CLEM experiments, duodenal monolayer cultures were generated from 28-week-old male Tph1-CFP mice. For quantitative RT-PCR of isolated EC cells, 8-12 weeks old wild type C57BL/6J male mice were used.

### Generation of Tph1-iCreERT2 knock-in mice

Tph1-iCreERT2 KI mice were generated using CRISPR-Cas9 gene editing technology similar to a previously published protocol^117^. Briefly, an HDR fragment was designed to replace the endogenous Stop codon of the mouse *Tph1* gene (Gene ID: 21990) by sequences encoding a GGAAGCGGA-Linker, a P2A self-cleaving peptide, a codon-improved Cre-recombinase (iCre)^118^, a GGAGGCAGTGGAGGCTCTGGAGGGTCT-Linker, and an estrogen receptor targeting motif (ERT2), and a Stop Codon (**Supplementary Item S1**). The oligonucleotide sequences used for the generation of the HDR fragment are listed in **Supplementary Table S2**. Mouse *Tph1* gene sequences [genomic C57BL/6J (Janvier) DNA; left arm, oligonucleotides #38360 and #38361; right arm, #38367 and #38368], the iCre [#38363 and #38364] and the ERT2 [#38365 and #38366] sequences were generated by PCR. The GGAAGCGGA-Linker and the P2A self-cleaving peptide were generated by PCR extension (#38346 and #38347). The PCR products were subcloned into pMiniT (NEB) using Gibson assembly^119^. hCas9-mRNA (25 ng/µl) was injected in combination with Cas9/sgRNA ribonuclear complexes [RNP; Cas9-Protein, 20 ng/µl; sgRNAs, 25ng/µl each; protospacer sequence of sgRNAs: *sgRNA1*, 5’-ATGCCCTCGCTAGGGTCACC-3’ (PAM = AGG); *sgRNA2*, 5’-CCACCTGGTGACCCTAGCGA-3’ (PAM = GGG)] into zygotes. The correct insertion of the HDR fragment in the offspring was tested by PCR using two primer pairs [#1 5’-TTGAGAGCTCCTGACTCATCCG-3 and 5’-CACAGGGAGGGCAGGCAGGTTTTG-3’ (1614 bp); #2 5’-TTCCTTGCAAAAGTATTACATCACGG-3’ and 5’-GGATTACACTTCCTTGTAGAGCA-3’ (1648 bp)]. We obtained one homozygous male Founder for which the correct insertion was confirmed by sequencing and, which was then crossed with C57BL/6J females to obtain germline-transmission of the knockin allele. Genotyping of this line is done with three primers [5’-TCCTGTGTTTAGTTTTGTCACCC-3’; 5’-TTCCTTGCAAAAGTATTACATCACGG-3’; 5’-AGAAGTAGAAGATAAGCAGTTCCA-3’; WT, 432 bp; KI, 248 bp]. The Tph1-P2A-iCreERT2 allele was further crossed into Ai32 animals^116^, which conditionally express a fusion protein of the light-gated ion channel Channelrhodopsin2 and EYFP from the *Rosa26* gene locus. 2D monolayer and 3D organoid cultures were generated from Tph1-iCreERT2;Ai32 mice to test for a potential “leakiness” of the iCreERT2 system by evaluating EYFP expression in the absence or presence of 4-hydroxytamoxifen (4-OHT), which induces nuclear translocation of iCreERT2 to induce Cre-mediated recombination.

### Cell lines

For the production of Wnt3a-and R-Spondin1-conditioned cell culture medium to supplement the 3D organoid culture medium, we followed previously published protocols^120, 121^. Cultrex HA-R-Spondin1-Tc 293T cells (Trevigen) were cultured in DMEM (high glucose, GlutaMAX, Gibco Cat# 31966021) supplemented with 10% FBS and 1% penicillin/streptomycin and 300 µg/ml Zeocin. When the cells reached ∼75% −80% confluence, the culture medium was replaced with organoid basal medium [Advanced DMEM/F-12 (Gibco, Cat# 12634010) supplemented with organoid basal medium 1% penicillin/streptomycin, 1% GlutaMAX and 10 mM HEPES] and R-Spondin1-conditioned medium was harvested after a week. L-Wnt3a cells (^122^; kindly provided by Dr. Hans Clevers) were cultured in DMEM (high glucose, GlutaMAX, Gibco Cat# 31966021) supplemented with 10% FBS and 1% penicillin/streptomycin and 125 µg/ml Zeocin and Wnt3a-conditioned medium was harvested in DMEM (high glucose, GlutaMAX, Gibco Cat# 31966021) supplemented with 10% FBS and 1% penicillin/streptomycin.

### Acute 2-dimensional (2D) intestinal epithelium monolayer cultures

To generate small intestine 2D monolayer cultures, we adopted a previously published protocol^43, 44^. Briefly, adult Tph1-CFP transgenic mice were sacrificed by cervical dislocation. The mice were opened from the abdomen, the small intestine was removed and the proximal small intestine (duodenum; approx. first 5 cm after the pyloric sphincter) was excised and immersed in 20 ml ice-cold L-15 medium. The gut tissue was then transferred in 10 cm petri dish containing ice-cold PBS (Sigma-Aldrich, D8662; containing MgCl_2_ and CaCl_2_) and cleaned from associated adipose tissue and the mesentery using a forceps and scissors. Using a pair of fine forceps, the muscle tissue around the extracted gut region was peeled off. The gut was then opened longitudinally with scissors to expose the epithelial layer and to allow the gut tissue to flatten. Gut villi were scraped off using a glass slide and the tissue was chopped into smaller pieces (approx. 2 x 2 mm) using a surgical blade. The tissue pieces were transferred into 50 ml centrifuge tube containing 20 ml of ice-cold PBS and they were washed several times in PBS. Next, the tissue fragments were then transferred into a fresh 50 ml centrifuge tube containing 8 ml prewarmed digestion solution [DMEM, high glucose (Sigma-Aldrich, D6546) containing collagenase enzyme (0.375 mg/ml), 37 °C], and incubated at 37 °C. After 5 minutes, the tube was shaken ∼10-20 times and the tissue pieces were left few seconds to settle. The supernatant was collected into a new tube and the digest was continued with fresh digestion solution. Tissue pieces were shaken every 5 minutes and fractions were collected every 5-10 minutes and inspected under an inverted microscope for the presence of crypts. Fractions containing a high density of crypts were collected by centrifugation (100 x g for 3 minutes). The supernatant was discarded, and the pellet was resuspended in prewarmed (37 °C) 2D culture medium [DMEM, high glucose (Sigma-Aldrich, D6546) containing 10% FCS, 100 U/ml Penicillin, 0.1 mg/ml streptomycin, 2 mM GlutaMAX, and 10 *µ*m Y-27632 ROCK inhibitor]. To remove large pieces of tissue debris, crypts were passed through a 70 *µ*m cell strainer and collected in 2D culture medium and plated on pre-coated coverslips (electrophysiology/electrochemistry and immunocytochemistry) or sapphire discs (electron microscopy) as described below.

For electrophysiology experiments, sterile round 25 mm glass coverslips were placed into individual wells of a 6-well plate, coated with ∼300 µl drops of ice-cold coating mix [DMEM - high glucose (Sigma-Aldrich, D6546)] containing 2% Cultrex Reduced Growth Factor Basement Membrane Extract (RGF BME)], positioned in the center of the coverslip and placed at 37 °C, 5% CO_2_ for at least 30 minutes. Immediately prior to the plating of the crypts, the coating mix was removed by aspiration. Crypts were plated in 2D-culture medium by pipetting 300 µl of concentrated crypts onto the coated regions of the coverslips and the cell culture plate was placed into a cell culture incubator for an hour at 37 °C, 5% CO_2_ for the crypts to settle. Then, 2.5 ml of 2D culture medium was added per well.

For immunocytochemistry experiments, sterile round 12 mm coverslips were placed into individual wells of a 24-well plate and coated either with 250 µl coating mix for at least 30 minutes at 37 °C, 5% CO_2_ or with an alternative coating solution (17 mM acetic acid, 1 mg/ml collagen, 0.5 mg/ml poly-D-lysine). Concentrated crypts were suspended in an appropriate volume of 2D culture medium, 500 µl of this suspension was added to each well after removal of the coating mix, and the plate was carefully placed into a cell culture incubator at 37 °C, 5% CO_2_.

For electron microscopy experiments, sapphire discs (Leica; #16770158) were prepared for cell culture experiments as previously described^123^. Briefly, discs were washed and sonicated in water and then again in pure ethanol. A thin (approximately 4 nm) carbon-coordinate system was deposited onto each disc using a CCU-010 HV vacuum coating unit (Safematic) and Pyser-SGI 135 mesh finder grids with an H15 reference pattern (Science Services; #PY135HF15-CR). Carbon-coated discs were baked for 12 h at 120 °C, and then stored at room temperature until use. On the culture day, discs were briefly sterilized by UV-exposure in a tissue culture hood and fixed with a small drop (∼1 µl) of Matrigel onto the glass bottom of a Nunc^TM^ Lab-Tek^TM^ coverglass chamber (Cat. No. 155380; 4 sapphire discs per well of a 2-well chamber). The Matrigel was allowed to polymerize for 30 minutes at 37 °C and the discs were then coated with small drops (∼30 µl) of coating mix for at least 30 minutes at 37 °C, 5% CO_2_. The coating mix was then aspirated, small drops (∼20 µl) of 2D culture medium containing concentrated crypts were plated onto each sapphire disc, the crypts were allowed to settle for 1 h at 37 °C, 5% CO_2_, and then approximately 2 ml 2D culture medium was added to each well.

To induce iCreERT2 translocation to the nucleus in cultures from the Tph1-iCreERT2;Ai32 mouse line, colonic cultures were generated using a protocol similar to the one described above for the small intestine and treated for 16 h on DIV1 with 1.25 *µ*m 4-OHT in the culture medium. The medium was subsequently changed to remove the 4-OHT, and cells were fixed and prepared for immunocytochemistry after an additional 8 h to allow more time for the expression of the fluorescent EYFP reporter.

### Small intestine 3D organoid cultures

Adult Tph1-CFP transgenic mice were sacrificed by cervical dislocation and small intestine organoid cultures were prepared according to previously published protocols^124–126^. Briefly, proximal small intestine (duodenum; approx. first 5 cm after the pyloric sphincter) was prepared as described for the 2D monolayer cultures, but without removing the muscle layer. The gut tissue was chopped in small pieces (approx. 2 x 2 mm) using a surgical blade. The tissue pieces were then transferred to a 50 ml tube containing 20 ml ice-cold PBS without MgCl_2_ and CaCl_2_ and washed extensively several times. The PBS of the final washing step was then discarded, and the tissue was treated in PBS supplemented with 2 mM EDTA for 30 minutes under gentle continuous rotation at 4 °C. The supernatant was then discarded, and the crypts were extracted from the intestinal tissue through trituration in ice-cold PBS. The supernatant containing the isolated crypts was collected. The trituration step was repeated 4 times and only fractions with a high density of crypts and few single cells were collected. To remove large pieces of tissue debris, the crypts were passed through a 70 *µ*m cell strainer. Crypts were collected by centrifugation (150 x g for 5 minutes) and the supernatant was completely removed. The crypts were resuspended in a small volume (approx. 20 µl) of prewarmed (at 37 °C) organoid basal culture medium [DMEM/F12, % GlutaMAX, 1% Penicillin/Streptomycin, 10 mM HEPES]. The resuspended crypts were then transferred into an aliquot of 200 µl of ice-cold Matrigel (Corning, Cat. #354230) and drops of 20 µl were placed carefully into individual wells of a prewarmed 48-well plate using precooled pipette tips. The Matrigel was allowed to polymerize for 15 minutes at 37 °C and 5% CO_2_ and 250 µl small intestine organoid culture medium [ENR; Advanced DMEM/F12 supplemented with 1 mM N-acetylcysteine, 10% R-Spondin1-conditioned medium, 5 µl of 100 ng/ml Noggin, 50 ng/ml EGF, 1x N2 supplement, 1x B27 supplement, and 10 *µ*m Y-27632] was then carefully added to each well. Note that all plastic tubes and pipettes / pipette tips except for the ones used to handle Matrigel were pre-coated with 100% fetal bovine serum (FBS). Crypts were cultured at 37 °C and 5% CO_2_ and medium was changed every 2-3 days.

Every 7-10 days, organoids were passaged to propagate the organoid line. Briefly, culture medium was aspirated and 500 µl of ice-cold basal culture medium was added to each well. The organoids were collected from individual wells and pooled in a 15 ml centrifuge tube. The organoids were released from the Matrigel by trituration. Organoids were harvested by centrifugation (150 x g for 5 minutes), and the supernatant including the Matrigel was discarded. Organoids were briefly digested in 0.5 ml warm (37 °C) TrypLE Express (Invitrogen, 12605010) and the digestion was immediately stopped by adding 0.6 ml Advanced DMEM/F12 medium supplemented with 10% FBS and 10 *µ*m Y-27632. The organoids were briefly dissociated by trituration and centrifuged at 150 x g for 5 minutes. The supernatant was discarded, and the pellet was transferred into an aliquot of ice-cold Matrigel and 20 µl domes were plated onto a prewarmed 48-well plate. Two to three days prior to experiments (biochemistry or plating as 2D cultures for electrophysiology), mature organoids were collected and embedded in fresh Matrigel.

To stimulate EC cell differentiation in mouse organoids using the *N*-cyclopropyl-5-(thiophen-2-yl)-isoxazole-3-carboxamide (ISX-9) pulse protocol^69^, crypts from the small intestine of a Tph1-CFP mouse were isolated and plated in 20 µl domes of Matrigel per well of a 48-well plate. The Matrigel was polymerized for 15 min at 37 °C and 250 µl stem cell growth medium containing Wnt3a-conditioned medium [WENR; Advanced DMEM/F12 supplemented with 1 mM N-acetylcysteine, 50% Wnt3A conditioned medium, 10% R-Spondin1-conditioned medium, 5 µl of 100 ng/ml Noggin, 50 ng/ml EGF, 1x N2 supplement, 1x B27 supplement, and 10 *µ*m Y-27632] was added. Three days later, medium was changed into ENR culture medium without Wnt3A and Y-27632, but supplemented with either 40 *µ*m ISX-9 dissolved in DMSO or with DMSO alone (vehicle control). After 48 h, the medium was removed and changed to ENR culture medium without ISX-9. ISX-9 or DMSO treated organoids were harvested for Western Blot experiments. To induce translocation of Cre-recombinase to the nucleus in the Tph1-p2a-iCreERT2;Ai32 line, duodenal organoids were embedded on DIV4 in fresh Matrigel and treated for 15 h with 1.25 *µ*m 4-OHT added to the organoid medium. Organoids were fixed and processed for immunofluorescence imaging 5 days later.

### Plating organoids as 2D-monolayer cultures

For functional experiments, organoids were converted to 2D monolayer cultures by adapting a previously published protocol^124^. Briefly, sterile round 25 mm glass coverslips were placed into individual wells of a 6-well plate, coated with ∼300 µl drops of ice-cold coating mix [DMEM - high glucose (Sigma-Aldrich, D6546)] containing 2% Cultrex RGF BME], positioned in the center of the coverslip and placed at 37 °C, 5% CO_2_ for at least 30 minutes. For plating, 2 wells of organoids were transferred on a single coverslip. Briefly, organoid culture medium was aspirated and 500 µl of ice-cold basal culture medium was added to each well. The organoids were collected from individual wells and pooled in a 15 ml centrifuge tube. The organoids were released from the Matrigel by trituration. Organoids were harvested by centrifugation (150 x g for 5 minutes), and the supernatant including the Matrigel was discarded. Organoids were briefly digested in 0.5 ml warm (37 °C) TrypLE Express and the digestion was immediately stopped by adding 0.6 ml Advanced DMEM/F12 medium supplemented with 10% FBS and 10 *µ*m Y-27632. The organoids were then briefly dissociated by trituration and centrifuged at 150 x g for 5 minutes. Immediately prior to the plating, the coating mix was removed from the coverslips by aspiration and the organoid fragments were plated in 2D culture medium [DMEM, high glucose (Sigma-Aldrich, D6546) containing 10% FCS, 100 U/ml Penicillin, 0.1 mg/ml streptomycin, 2 mM GlutaMAX, and 10 *µ*m Y-27632] by pipetting 300 µl of the cell suspension onto the coated regions of the coverslips. The cell culture plate was then placed into a cell culture incubator for an hour at 37 °C, 5% CO_2_ for the crypts to settle. Then, 2.5 ml of 2D culture medium was added per well. The cultures were used after 24 h for functional experiments.

### Generation and culture of human small intestine organoids

The generation and culturing of human terminal ileum organoids was described elsewhere^69^. Briefly, human organoids were cultured in Matrigel (Corning, #356231) and culture medium [WENRAS; Advanced DMEM/F12 supplemented with 1 mM N-acetylcysteine, 50% Wnt3A conditioned medium, 10% R-Spondin1-conditioned medium, 5 µl of 100 ng/ml Noggin, 50 ng/ml EGF, 1x N2 supplement, 1x B27 supplement, 10 nM gastrin, 500 nM A83-01, 10 *µ*m SB202190, 10 mM nicotinamide, and 10 *µ*m Y-27632]. For the differentiation of human organoids, Wnt3a conditioned medium was reduced from 50% to 15%, SB202190 and nicotinamide were withdrawn, and five days after passaging organoids were treated with 40 *µ*m ISX-9 for 48 h. Afterwards, ISX-9 was removed for another 48 h so that organoids were harvested for Western Blot experiments 96 h after the beginning of the ISX-9 pulse.

### Adrenal chromaffin cell cultures

Adrenal glands were extracted from adult mice of either gender following standard protocols^50, 127^. Adult animals were sacrificed before the extraction by cervical dislocation. The animal was then cut open at the abdomen and the adrenal glands were extracted from the surrounding connective tissue and placed on ice-cold ringer-Locke’s solution [0.15 M NaCl, 5.6 mM KCl, 0.85 mM NaH_2_PO_4_ (anhydrous), 2.15 mM Na_2_HPO_4_X (H2O), 10 mM D-Glucose]. The glands were then cleaned from the connective tissues. Adrenal chromaffin cell culture medium [DMEM (Linaris, GMF2143YK) containing 0.2% Penicillin-Streptomycin and 1% ITSX] was prepared freshly. In parallel, papain-enzyme solution [DMEM supplemented with 2 mg/ml L-Cysteine, 1 mM CaCl_2_, 0.5 mM EDTA] was prepared by adding 20 units of papain extract per 1 ml of enzyme solution and equilibrating the solution by carbogen bubbling for 10 minutes. The pair of glands was washed shortly in a drop of enzyme solution and then incubated in 300 µl enzyme solution under continuous shaking at 37 °C for 45 minutes. The enzyme solution was then completely removed, replaced with 300 µl of inactivation solution [DMEM supplemented with 2.5 mg/ml bovine serum albumin, 2.5 mg/ml Trypsin inhibitor, 10% (v/v) FBS] and incubated for 5 minutes at 37 °C. The inactivation solution was then completely removed, and the adrenal glands were washed in 500 µl adrenal chromaffin cell culture medium. The medium was removed, 200 µl fresh culture medium was added, and the gland was triturated to release the chromaffin cells from the adrenal medulla. An additional 100 µl of medium were added to reach a final volume of approx. 300 µl, and the cells were then seeded onto uncoated, round, 25 mm glass coverslips in a 6-well plate as drops of 50 µl. The cells were allowed to settle and adhere on the glass coverslip for 30 minutes in a cell culture incubator at 37 °C, 5% CO_2_. Then, 2 ml of DMEM was carefully added to each coverslip and the isolated cells were cultured at 37 °C, 5% CO_2_. Electrophysiological recordings were performed after 48 h.

## Method details

### Immunocytochemistry

For immunocytochemical analyses of EC cells in 2D monolayer cultures, coverslips were washed 24 h after plating three times with PBS and cells were fixed in ice-cold fixative [4% (w/v) paraformaldehyde (PFA) in 0.1 M phosphate buffer (PB), pH 7.4] for 30 minutes at room temperature. Coverslips were then washed three times in 0.1 M PB, incubated for 45 minutes at room temperature in blocking and permeabilization solution [0.1 M PB, 10% (v/v) goat or donkey serum, 0.3% (v/v) Triton-X-100, 0.1% (w/v) fish skin gelatine], incubated for 2 h in incubation buffer [0.1 M PB, 5% (v/v) goat or donkey serum, 0.1% (v/v) Triton-X-100] containing primary antibodies, washed several times in 0.1 M PB, and incubated in secondary antibodies in incubation solution. Cells on coverslips were then washed several times in 0.1 M PB and stained in 300 nM DAPI for 10 minutes at room temperature (Sigma-Aldrich; #10236276001) to label cell nuclei. Following a couple of brief washes in 0.1 M PB, slides were dipped in distilled water and coverslips were mounted using Aqua-Poly/Mount mounting medium (Polysciences, Inc.; #18606-20) onto glass slides for light microscopic analysis. The following primary antibodies were used: rabbit polyclonal anti-CAPS1 (Synaptic Systems #262 003, dilution 1:500), rabbit polyclonal anti-Rab3 (Synaptic Systems #107 003, dilution 1:500), rabbit polyclonal anti-Secretin (Phoenix Pharmaceuticals #H-067-04, dilution 1:2,000), guinea pig polyclonal anti-Substance-P (Abcam #ab10353, dilution 1:200), goat polyclonal anti-5-HT (Immunostar #20079, dilution 1:2,000), rabbit polyclonal anti-5-HT (Immunostar #20080, dilution 1:2,000) and rabbit polyclonal anti-GFP (MBL #MBL-598, dilution 1:500:). The following secondary antibodies were used: donkey polyclonal anti-rabbit Alexa555 (Themo Fisher #A631572, dilution 1:2,000), goat polyclonal anti-rabbit Alexa488 (Thermo Fisher #A11008, dilution 1:2,000), goat polyclonal anti-guinea pig Alexa555 (Thermo Fisher #A21435, dilution 1:2,000) and donkey polyclonal anti-goat Alexa633 (Mobitec #21082, dilution 1:2,000).

Mature mouse organoids in Matrigel were washed three times with PBS and fixed in ice-cold fixative [4% (w/v) paraformaldehyde (PFA) in 0.1 M phosphate buffer (PB), pH 7.4] for 30 min at room temperature. Immunofluorescence staining of organoids was performed as described above. Organoids were then transferred carefully with a brush onto a glass slide and mounted in Aqua-Poly/Mount mounting medium.

#### Immunohistochemistry

For immunohistochemical characterisations of EC cells in gut tissue, animals were sacrificed by cervical dislocation and the small intestine was removed from the body. The proximal small intestine (duodenum; approx. first 5 cm after the pyloric sphincter) was cut into small (approx. 1 cm) long tissue pieces using scissors, which were immersed in ice-cold fixative [4% PFA in 0.1 M PB (pH 7.4)]. Gut tissue pieces were fixed for 2 hours at 4 °C, then washed several times in 0.1M PB and cryoprotected by immersion in an increasing gradient of 20% and 30% sucrose in 0.1 M PB (pH 7.4). Gut tissue pieces were then briefly immersed in liquid Tissue-Tek^®^ OCT compound (Sakura, #4583) and several gut pieces were aligned next to each other in an aluminium foil embedding form partially filled with precooled OCT compound. The aluminium form containing the tissue pieces was then fully filled with liquid OCT, the sample was rapidly frozen on a liquid nitrogen-cooled aluminium block and stored at −80°C. One hour before cryo-sectioning, the OCT block was transferred to a −20°C freezer and then removed from the aluminium foil and mounted on a cryostat (Leica CM3050 S) stub. Cross-sections (10 *µ*m-thick) of the intestinal tissue pieces were cut and the cryo-sections were thaw-mounted on Superfrost™ slides. The tissue slides were then washed briefly in 0.1 M PB (pH 7.4), incubated for 1 h at RT in blocking solution [5% goat or donkey serum, 0.3% Triton X-100, and 0.1% cold-water fish skin gelatine in 0.1 M PB (7.4) overnight], and then incubated in primary antibodies diluted in 3% goat or donkey serum, 0.1% Triton X-100, and 0.1% fish skin gelatine in 0.1 M PB (pH 7.4). The primary antibodies used for immunohistochemistry included: rabbit polyclonal anti-CAPS1 (Synaptic Systems #262 003, dilution 1:500), and rabbit polyclonal anti-Rab3 (Synaptic Systems #107 003, dilution 1:500). Tissue sections were then washed in 0.1 M PB (pH 7.4) and incubated in fluorophore-conjugated secondary antibodies diluted in 3% goat or donkey serum, 0.1% Triton X-100 and 0.1% fish skin gelatine in 0.1 M PB (pH 7.4) for 2 h at room temperature. The secondary antibody used in the study was donkey polyclonal anti-rabbit Alexa555 (Themo Fisher #A631572, dilution 1:2,000). Tissue sections were then washed in 0.1M PB and incubated for 30 min at RT in 300 nM DAPI (Sigma-Aldrich; #10236276001) to label cell nuclei. Following a couple of brief washes in 0.1 M PB, slides were dipped in distilled water and Menzel-Gläser #1.5 coverslips were mounted using Aqua-Poly/Mount mounting medium (Polysciences, Inc.; #18606-20).

To visualise endogenous CFP-fluorescence in gut cross-sections of the Tph1-CFP mouse line, tissue pieces of the duodenum and colon were dissected and immersion-fixed with ice-cold fixative [4% PFA in 0.1 M PB (pH 7.4)] as described above. After the fixation, the samples were washed in 0.1M PB, embedded in agarose and 50 *µ*m-thick sections were cut on a VT1200 S vibrating microtome (Leica). Slices were mounted onto glass slides using Menzel-Gläser #1.5 coverslips and Aqua-Poly/Mount mounting medium (Polysciences, Inc.; #18606-20) and imaged without further processing.

### Light microscopic analysis of immunolabelled EC cells

The visualization of immunolabelled EC cells in monolayer cultures from Tph1-CFP mice was achieved using a widefield microscope (Leica DMi8, Thunder 3D Live Cell Imaging System) equipped with a Spectra X light engine (Lumencor), an LMT200 mechanical high-precision mechanical stage (ITK), and a 4.2 MP DFC9000 sCMOS camera (Leica). Tiled DIC and fluorescent multi-channel images were acquired through 20x (HC PL FL L 20x/0.40 CORR PH1; Leica) and 40x (HC PL APO 40x/0.95 CORR PH2; Leica) magnification objectives, respectively. Based on DIC overviews, regions with a good coverage of cells were selected for fluorescent imaging at 40x magnification (16-bit; unbinned pixel sizes x, y, 0.163 *µ*m). Different combinations of LED excitation light and fluorescent filter cubes were used to visualize fluorescent secondary antisera: DAPI (395 nm excitation, DFT51011 filter set), CFP (440 nm excitation, CFP filter cube), Alexa488 (470 nm excitation, DFT51011 filter set), Alexa555 (550 nm excitation, DFT51011 filter set) and Alexa633 (640 nm excitation, DFT51011 filter set). Z-stacks were acquired and computationally cleared using Leica Acquisition Suite X (LAS X; Leica) and patented THUNDER technology (Leica). Identical parameters were applied for all quantified images (image acquisition: z-depth, step size; image processing: feature scale, regularization, iterations). All data were saved as LIF files until further processing.

### Image analysis

To test whether all serotonergic cells in intestinal 2D epithelial monolayer cultures from the Tph1-CFP mouse line express the fluorescent reporter, duodenal and colonic cultures were labeled with an antibody directed against GFP to enhance endogenous CFP-fluorescence, an antibody detecting 5-HT to detect all serotonergic cells in culture, and DAPI to visualize all cell nuclei. The specific goal was to quantify the number of 5-HT-immunopositive cells that were positive for Tph1-CFP and vice versa, the number of Tph1-CFP positive cells that are positive for 5-HT. Light microscopic images were analysed in FIJI with custom-written ImageJ macros (ImageJ 2.1.0/1.53h)^128^.

Firstly, image stacks of the four respective channels (DAPI, Tph1-CFP, anti-GFP, anti-5-HT) were imported into FIJI, z-projected using the “Sum Slices” function and exported as TIF files. A threshold that was initially determined manually to detect all DAPI positive nuclei was then applied on the created TIF files in the DAPI channels. A watershed algorithm was used to separate overlapping signals. The number of DAPI positive nuclei per image was then determined with the “Analyze Particles” function.

Secondly, regions of interest (ROIs) were created for the anti-GFP and anti-5-HT channels, respectively. For this, a threshold that includes positive cells but not cell debris was determined manually once, and this threshold was then applied to all images. The “Analyze Particles” routine was then used to create the actual ROIs. In the next step, these ROIs were used to measure the signal intensity in the respective other channels. Following that protocol, ROIs were created in the anti-GFP channel to measure the individual signal intensities in the Tph1-CFP channel. Likewise, signal intensities in the ROIs from the 5-HT channel were measured in the Tph1-CFP and the anti-GFP channels. Apart from the mean signal intensity following parameters were additionally quantified: intensity minimum, maximum, integrated density and modal values with the corresponding standard deviation of the mean as well as the ROI area, perimeter, shape descriptors, Feret’s diameter and area fraction. These data were exported as CSV files and systematically imported into Excel (Microsoft Corporation) using a custom macro written in Visual Basic for Applications (VBA). For the analysis of these data, a threshold was calculated for which a signal intensity was considered above background. To achieve this, the intensity outside the ROIs was measured per image and the value corresponding to two standard deviations above the mean intensity of all images was chosen.

### Live cell imaging for correlative light electron microscopy (CLEM)

Live imaging of 2D monolayer cultures on sapphire discs was performed to map the locations of CFP-positive EC cells using a widefield microscope (Leica DMi8, Thunder 3D Live Cell Imaging System) equipped with a Spectra X light engine (Lumencor), an LMT200 mechanical high-precision mechanical stage (ITK), and environmental control (Large Incubator, Heating Unit 2000, TempController 2000-2, CO_2_-Controller 2000, CO_2_-Cover; Pecon). Cultured cells were maintained at 37 °C and 5% CO_2_ levels at all times during image acquisition. Tiled, multi-channel images (16-bit; pixel sizes x, y, 0.325 *µ*m) encompassing the deposited carbon coordinate system were acquired through an HC PL APO 40x/0.95 objective (Leica; #11506415) with a 4.2 MP DFC9000 sCMOS camera (Leica). The positions of CFP-expressing EC cells were visualized using 440 nm excitation in combination with a CFP filter cube (Leica; #11525305) and related to the generalized pattern of all cultured cells and the carbon coordinate system revealed by DIC optics. LAS X software (Leica) was used to implement an interpolated focus strategy, which provided an optimal focus depth at all imaged positions of the sapphire disc, and to merge tiled images. Signal intensities and contrast were adjusted using Fiji software^128^.

### High-pressure freezing

Immediately after live imaging, cultured 2D monolayers were rapidly cryofixed with an EM ICE high-pressure freezing device (Leica). Individual sapphire discs were carefully removed from the underlying glass of the LabTek chamber and transferred into HEPES-buffered extracellular solution kept at room temperature [(in mM): 145 NaCl, 2.8 KCl, 2 CaCl_2_, 1 MgCl_2_, 10 HEPES, and 11 D(+)glucose; pH was adjusted to 7.2 using NaOH and the osmolarity to 310 mOsm/kg using Mannitol] in a plastic petri dish used for preparation of the sapphire disc freezing assembly. A mylar spacer ring (outer diameter, inner diameter, depth = 6 x 5 x 0.1 mm, Leica; #16771883) was carefully positioned on top of the disc carrying the cultured cells and a new (uncoated) sapphire disc was placed on top of this assembly to serve as the “lid”. This assembly step was performed while all three components were fully submerged in the extracellular solution to prevent air bubbles from being trapped between the discs. The sapphire disc assembly was then transferred to a sample holder middle plate (Ø 6.0 mm; Leica; #16771838) and covered with a rubber cover ring (500 *µ*m; Leica; #16771884). Excess liquid was removed with Whatman filter paper and the middle plate with the sample was positioned on top of a half cylinder (Leica; #16771846) on the stage of the HPF device. High-pressure frozen sapphire discs subsequently stored in liquid nitrogen until further processing by automated freeze-substitution.

### Automated freeze substitution

High-pressure frozen cultures on sapphire disc freezing substrates were loaded into the AFS2 freeze substitution device (Leica) and cryo-substituted as previously described ^123^. Cryo-fixed samples were removed from liquid nitrogen storage and immersed in anhydrous acetone precooled to −90 °C. The sapphire disc assembly was carefully separated and the sapphire disc bearing cells was carefully loaded with cryo-forceps into a custom-built aluminium sapphire disc revolver^123^ immersed in 0.1% tannic acid in anhydrous acetone at −90 °C. Automated freeze substitution was performed as previously described^129, 130^ with minor modifications. Briefly, samples were incubated in 0.1% tannic acid in anhydrous acetone for 2 days at −90 ^°^C, fixed and metal stained with 2% osmium tetroxide in anhydrous acetone as the temperature slowly ramped up over several days to 4 °C, washed in acetone and brought to room temperature for EPON resin infiltration and embedding. For the embedding, sapphire discs were placed slice-upwards on Parafilm-coated glass slides, gel-capsules filled with 100% EPON were inverted on the specimen, and the samples were polymerized at 60 ^°^C for 24-48 h. Polymerized blocks were removed from the glass slides and the sapphire discs were carefully trimmed off using a razor blade and brief contact with a liquid nitrogen-cooled cotton swab, thus exposing the carbon-coordinate system on the block-face. To define an appropriate region of interest (R.O.I.) for subsequent ultramicrotomy, the distribution of CFP-expressing EC cells determined by live cell imaging was related to their corresponding carbon coordinates. An EM TRIM 2 high-speed milling device (Leica) equipped with a diamond bit was used to refine the block-face to the selected R.O.I. by visualizing the carbon coordinate system in reflected light. The dimensions of the block-face R.O.I. (approximately 1.1 x 0.4 mm) were designed to permit multiple serial sections to be positioned and viewed in their entirety on a 3 mm electron microscopy grid.

### Ultramicrotomy

A DiATOME histo diamond knife (Science Services; #DH4560) mounted on an EM UC7 ultramicrotome (Leica) was used to cut sections from EPON-embedded slices according to the following sequence spanning a cutting depth of approximately 2 *µ*m: 4 x 200 nm, 3 x 60 nm, and 3 x 350 nm. This sequence was repeated, and the last 350 nm-thick section of each sequence was collected on a glass slide and Nissl stained to assess the structural organization of the monolayer culture. Ultrathin (60 nm) and semithin (200 nm and 350 nm) sections were collected in series on Formvar-filmed, carbon-coated, and glow-discharged copper grids for 2D transmission electron microscopy (Gilder parallel 75 mesh; Science Services; #G75PB-CU) and 3D electron tomography (Gilder hexagonal 100 mesh; Gilder; #G100H-CU), respectively. Ultrathin sections were contrasted with 1% aqueous uranyl acetate and 0.3% Reynold’s lead citrate. Semithin sections were treated with Protein A coupled to 15 nm or 10 nm gold particles (Cell Microscopy Center, Utrecht; The Netherlands), which served as fiducial markers for tilt-series acquired at 11,000 x and 22,000 x magnification, respectively. Prior to electron tomographic analysis, the quality of cryo-preservation was assessed in ultrathin sections according to previously established criteria^131^.

### Electron tomography and data analysis

To resolve the 3D ultrastructural organization of EC cells and quantify the polydispersity of secretory granules, transmission electron tomography was performed using a Talos F200C G2 scanning/transmission electron microscope (Thermo Scientific) equipped with an X-FEG operating at 200 kV acceleration voltage. Tiled images of serial sections on EM grids were acquired at two magnifications using MAPS 3 software (Thermo Scientific): (i) an overview of section coordinates on the grid was obtained at 84 x magnification; and (ii) the positions of individual cells within the borders of respective plastic sections were mapped in tiled images acquired at 5000 x magnification. The latter cell position maps were exported as merged tif images and imported together with corresponding images of the same R.O.I. acquired by live cell imaging into Icy software^132^. Correlative light-electron microscopy (CLEM) registration was performed at two levels of stringency using the ec-CLEM plug-in^133^. Registration was initially performed using a non-rigid transformation to align the carbon coordinate boundaries of the R.O.I. resolved in DIC images acquired during live cell imaging with the corners of plastic sections visualized in the electron microscope. Transformations applied to both DIC and CFP channels and typically indicated mild compression of plastic sections in the cutting direction. Upon import of the transformed light microscopic data into MAPS software and its alignment with individual plastic sections, the position of CFP-expressing cells could be determined with respect to the microscope stage coordinates and cells harbouring dense-core secretory granules could be reliably localized and visualized within close vicinity of light microscopic predictions (i.e. CFP fluorescence). The microscope stage positions of a few broadly distributed and isolated candidate cells thus identified by this approach were subsequently used as more stringent registration landmarks to further refine the prediction of cell positions to within < 5 um. This approach permitted not only the ultrastructural organization of CFP-expressing cells to be determined, but also that of non-serotonergic EECs for which no corresponding CFP signal was detected.

Single-axis tilt series of identified CFP-expressing EC cells were collected from −60 to +60 in 1° increments at 11000 x or 22000 x (pixel size x,y = 0.66 nm) magnification using TEM Tomography 4.x data acquisition software (Thermo Scientific) and a CETA 16M CMOS camera (Thermo Scientific) (pixel sizes x,y = 1.35 nm or 0.66 nm for 11000 x and 22000 x magnification, respectively). Tilt series were reconstructed into 3D volumes by weighted back-projection (binning = 2; voxel dimensions x,y,z = 1.32 nm or 2.70 nm for 11000 x and 22000 x magnification, respectively) and vesicular organelles were segmented for morphometric analysis using the ETOMO and 3dmod components of the IMOD software package^134^, respectively.

Reconstructed tomograms were segmented using the IMOD package^134^ (Kremer et al., 1996) similar to the method described previously^135^: All vesicles were segmented manually as perfect spheres with the center being placed into the tomographic slice, in which the vesicle diameter appeared to be the largest (i.e., when the vesicle is cut at its midline). For vesicles that did not appear as perfect spheres in the tomogram, the sphere size was adjusted with a similar overlap inside and outside the vesicle to get a good approximation of the “real” diameter. Vesicles that exhibited a prominent electron-dense core were segmented as LDCVs and vesicles with a clear core as small clear-core vesicles (SCCVs). The radius of each vesicle was extracted using the “imodinfo” command and imported into Excel using a custom-written VBA macro.

### Fabrication of carbon fibres

Carbon fibres (5 *µ*m diameter from goodfellow) were spread on a glass plate. Single fibres were picked and glued to high quality ethanol-cleaned copper wires using silver conductive. The fibre attached to the copper wire was then carefully inserted into a glass pipette (1.5 OD, Harvard apparatus GC150T). The pipette tip at the copper wire side was then glued using two-component epoxy glue and the glue was left to dry for at least two hours. Then, the pipettes were pulled near the connection point between the carbon fibre and the copper wire. The carbon fibre protruding the pipette tip was then electroplated with electrodeposition paint (EDP) at 5V for about 40 seconds. EDP consisted of 15% of a 1:4 cymel 303 acrylic resin / viacryl SSC 6280w/45WA mixture diluted in distilled water. The mix was left to homogenize under gentle stirring overnight and stored at room temperature until use. Each microelectrode was observed individually under the microscope to make sure that a thin layer of the anodic EDP was covering the exposed part of the fibre. The microelectrodes were heat treated for 30 minutes at 170 °C. In cases where the tip of the pipette was not properly sealed, this was achieved using Sylgard 184 (Sigma, 761036) curing agent in combination with a heat-treatment using a hot air blower for about 60 seconds at medium ventilation. This procedure helped to achieve a better signal-to-noise ratio in the amperometric recordings by preventing extracellular solution leaking into the glass pipette.

### Electrochemistry

For functional studies of EC cells, electrophysiological recordings were performed either 24 h post-culture (acute 2D cultures) or 24 h after plating organoids as 2D monolayers. To gain easier access with the patch-pipette to the individual EC cells, each coverslip was incubated for 1 minute in prewarmed (37 °C) 0.05% TrypLE solution. Coverslips carrying cultured adrenal chromaffin cells were directly transferred into the recording chamber. The extracellular solution for both sets of experiments was composed of (in mM): 145 NaCl, 2.8 KCl, 2 CaCl_2_, 1 MgCl_2_, 10 HEPES, and 11 D(+)glucose; pH was adjusted to 7.2 using NaOH and the osmolarity to 310 mOsm/kg using Mannitol. All recordings were performed at room temperature. Cells were visualized with an inverted microscope (Zeiss Axiovert 200). In the case of EC cells, only cells demonstrating a prominent CFP-fluorescence were selected for functional studies and the experimental design was as follows: First, the carbon fibre microelectrode was lowered into the bath solution and moved to a position in the field of view near the cell. Next, positive pressure was applied to the patch pipette to prevent backflow of the extracellular solution when inserted into the bath solution. The pipette was lowered slowly onto the cell until touching it gently. The pressure was released to ease the formation of a tight gigaohm seal between the pipette and the cell plasma membrane. The series resistance (Rs) was automatically compensated to about 80%. The carbon fibre was then moved to gently touch the surface of the cell. Finally, gentle suction pulses were applied to the patch pipette to rupture the plasma membrane without disturbing the gigaseal and to infuse the cell with an intracellular solution containing a high concentration of free Ca^2+^ to trigger exocytosis. Amperometric currents were recorded from the carbon-fibre in close proximity to the cell plasma membrane immediately after opening the cell.

Patch pipettes were pulled from Capillary Glass (1.5 OD, Harvard Apparatus, GC150T-10) to have an open-tip resistance of 3.5 - 5.5 MΩ using a Sutter P-2000 horizontal puller. The tips of the pipettes were heat polished. The intracellular solution for “medium” calcium-infusion experiments was composed of (in mM): 100 Cs-glutamate, 6.67 BAPTA, 13.33 Ca-BAPTA, 2 Mg-ATP, 0.3 GTP, and 1 ascorbic acid, pH 7.2 (adjusted to 290 mOsm). The intracellular solution for “high” calcium-infusion experiments was composed of (in mM): 100 Cs-glutamate, 30.5 DPTA, 9.5 Ca-DPTA, 2 Mg-ATP, 0.3 GTP, and 1 ascorbic acid, pH 7.2 (adjusted to 290 mOsm). These intracellular solutions were based on solutions that are routinely used to calibrate the fluorescent signal ratios a 350/380 illumination for ratiometric calcium-imaging experiments, which however additionally contain the fluorescent calcium dyes Fura-4F and Furaptra^136, 137^. We estimated the free [Ca^2+^] for the “moderate” calcium-infusion experiments and “high” calcium-infusion experiments to be <5 *µ*m and > 20 *µ*m, respectively.

Amperometric recordings were performed as previously described^49, 138^ using 5 *µ*m carbon fibre microelectrodes. The carbon fibre microelectrodes were held at 650 and 800 mV relative to the ground electrode for EC and adrenal chromaffin cells, respectively, using a VA-10 amplifier (NPI electronic instruments). The amplifier was set to a gain of 5 mV/pA and the measured currents were digitized at 25 kHz and filtered online at 3 kHz. Exocytosis of vesicles was stimulated by infusing the cells with intracellular solutions containing high concentrations of free Ca^2+^ through the patch pipette.

### Analysis of amperometry measurements

The kinetic properties of the amperometric spikes were analysed in IgorPro (Wavemetrics) using a freely-available macro^48^. Currents were filtered off-line using a Gaussian filter with a cut-off at 3 kHz. Spike detection threshold was set to 5 pA and spikes that had more than 50% overlap were excluded from kinetic analyses, as it was impossible to fully fit the amperometric spike and estimate the half-width. Due to this exclusion criterion, we did not report the inter-spike interval (ISI), but quantified the total number of spikes detected throughout the 120s recording prior to applying the exclusion criterion mentioned above. Only spikes with a maximal amplitude of <6 pA and a charge of <6000 fC were included into the analysis and additionally spikes for which the 50-90% rise time couldn’t be fitted were excluded. For each spike, the median peak spike amplitude, spike charge, rise-time (rate by which the spike reaches 50% - 90% of its peak value), and half-width (width of the amperometric spike at the amplitude half-maximal value) were analysed. The prespike foot signal was determined manually for each spike. Cells with less than 10 spikes were excluded from the analysis^50^.

### Immunoblotting

Organoids in culture were either treated with ISX-9 or DMSO (vehicle control) and were harvested from the Matrigel. Organoids were washed several times in PBS, concentrated by centrifugation at low speed, flash-frozen in liquid nitrogen and stored at −80 °C until further use. On the day of the experiment, the organoid pellet was homogenized with a syringe in lysis buffer (320 mM sucrose, 5 mM HEPES pH 7.4, 1 mM EDTA, 0.1 mM PMSF, 0.5 µg/ml Leupeptin, 1 mg/ml aprotinin). The protein concentration was determined by the Bradford Assay and homogenates were dissolved in SDS buffer for sodium dodecyl sulphate polyacrylamide gel electrophoresis (SDS-PAGE) and heated for 5 minutes at 95 °C. Protein samples (20 µl/lane) were separated on Novex NUPage 4-12% Bis-Tris gradient gels in running buffer (50 mM MOPS, 50 mM Tris-Base, 1 mM EDTA, 0.1% (w/v) SDS, pH 7-7) and blotted overnight onto a nitrocellulose membrane in transfer buffer (25 mM Tris-Base, 190 mM glycine, 20% (v/v) methanol). After the transfer, protein bound to nitrocellulose membranes was visualized using the Pierce Reversible Protein Stain Kit for Nitrocellulose Membranes (MEM Code) assay following the manufacturers’ instructions. At this stage, the membranes were cut into smaller pieces, to be labelled with different antibodies. After MEM Code destaining, the membranes were washed with TBS buffer (20 mM Tris-Base, 137 mM NaCl, pH 7.5), incubated with blocking buffer (TBS, 5% (w/v) milk powder, 5% (v/v) goat serum, 0.1% (v/v) Tween-20) for 30 minutes, incubated with primary antibodies for 1 h at room temperature, washed in TBS-T buffer (TBS, 0.1% (v/v) Tween-20), and incubated with secondary antibodies conjugated to horseradish peroxidase. After several washing steps in TBS-T and TBS, immunoreactive bands were visualized with an enhanced chemiluminescence (ECL) detection system (Intas Science Imaging Instruments, Göttingen, Germany). The following primary antibodies were used: anti-Munc13-1 [#40, rabbit polyclonal, N-terminal, 1:1,000^139, 140^; anti-actin (Sigma-Aldrich #A470, mouse monoclonal, 1:2,000); anti-CAPS1 (SySy #262 003, rabbit polyclonal, 1:500), anti-Chromogranin A (SySy #259 003, rabbit polyclonal, 1:1,000); anti-GFP (MBL #598-7, rabbit polyclonal, 1:1,000); anti-Rab3 (SySy #107 003, rabbit polyclonal, 1:400); anti-SNAP25 (SySy #111 002, rabbit polyclonal, 1:2,000); anti-VAMP 2 (SySy #104 202, rabbit polyclonal, 1:400). The following secondary antibodies were used: Goat anti-Mouse IgG (H+L) and Goat anti-Rabbit IgG (H+L) coupled to HRP (Jackson ImmunoReseaarch, 1:10,000). Stained nitrocellulose membranes were scanned and served as a visual loading control.

### Quantitative RT-PCR of isolated EC cells

To obtain expression levels of various vesicle-associated proteins present in murine enterochromaffin cells, analysis were performed on RNA isolated in a previous in house study^36^. In short, the small intestine of standard chow diet fed, non-fasted, wild type C57BL/6-J male mice, 8 to 12 weeks old, and the proximal small intestine was dissected. The two segments were digested into a single cell suspension via collagenase treatment. The single cells were then fixed, gently permeabilized with saponin and the enterochromaffin cells were labeled intracellularly with a primary 5-HT (ab66047, Goat Anti 5-HT, 1:3200, Abcam) and Alexa488-conjugated secondary antibodies to enable FACS isolation. Cells were sorted into a 488 nm excitable fraction (5-HT positive) and a 488 nm non-excitable (5-HT negative) fraction using MoFLo Astrios (Beckman Coulter). Total RNA was isolated, and RT-PCR was performed using the SuperScript III Reverse Transcriptase (Invitrogen, Carlsbad, CA). Quantitative RT-PCR was run to characterise and confirm the isolated cell fractions with SYBR PrecisionPLUS (Primerdesign) and primers (Tph1, ChgA, Ywhaz, GAPDH from^141^) in a LightCycler480 (Roche). Due to the small sample size, cDNA was amplified with QuantiTect Whole Transcriptome kit (Qiagen). Custom-designed qPCR arrays for the expression of various presynaptic proteins and neuronal SNAREs (Qiagen) were used according to manufacturer’s instructions. Target regions are listed in **Supplementary Table S3**. To calculate relative expression we preceded as in^142^. Around 0.3% of total mucosal single cells were sorted as 5-HT positive EC cells from the small intestine. *Tph1* was on average enriched 1700 fold and *ChgA* 1400 fold in the small intestinal 5-HT positive cells (Fig. 1 in ^36^). Thus, the FACS-purified EC cells were very pure.

## Quantification and statistical analysis

### Statistical analysis

Data exploration and transformation for statistical analysis were performed in R using the tidv verse package. Statistical analyses were carried out using GraphPad Prism software 9 (* when p<0.05; ** when p<0.01, and *** when p<0.001). The data sets were tested for normality using the Kolmogorov-Smirnov test. If the data set was not normally distributed, a Mann-Whitney test was performed to probe for statistical significance. Normally distributed data sets were tested for statistical significance using unpaired two-tailed *t-*tests. Unless indicated otherwise, (n) refers to the number of tomograms (3D) analysed for each experiment, or the number of cells measured for electrophysiological experiments pooled over several cultures. (N) refers to the number of cultures / animals. Numbers of biological replications for each experiment are listed in the Figure legends and **Supplementary Table S1.** In the Figures, data are represented as mean ± SEM unless indicated otherwise. For electrophysiology experiments the median amplitude, charge, half-width, and rise time as well as the prespike foot amplitude, charge, and duration were calculated for each cell and the mean ± SEM of the medians were then plotted for data representation. Further, the average number of spikes per second per cell and the percentage of spikes with a prespike foot signal per cell were plotted as mean ± SEM. For the ultrastructural analysis of vesicle sizes, the mean vesicle diameter, vesicle volume per cell, and percentage of SCCVs and LDCVs from all vesicles analysed in a given tomogram were plotted. Plots were generated using either GraphPad Prism software 9 or the ggplot2 package of tidyverse and data visualization was carried out with the CorelDRAW software.

**Supplementary Table S1.** Experimental Animals

**Supplementary Table S2.** Oligonucleotides for the generation of the Tph1-P2-iCreERT2 mouse line

**Supplementary Table S3.** Target sequence of genes analysed by qPCR

**Supplementary Item S1.** Tph1-P2A-iCreERT2 Sequence Information

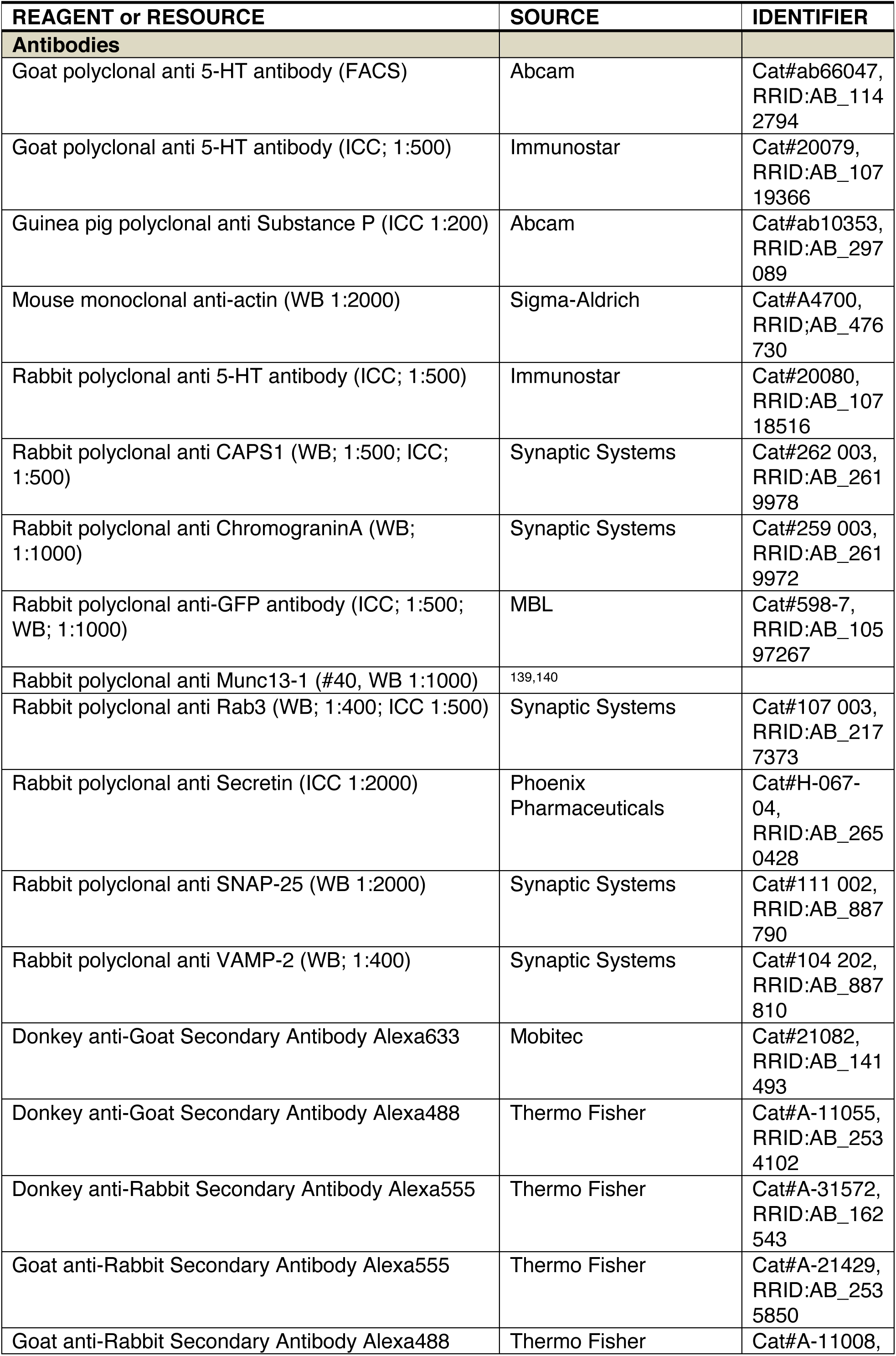
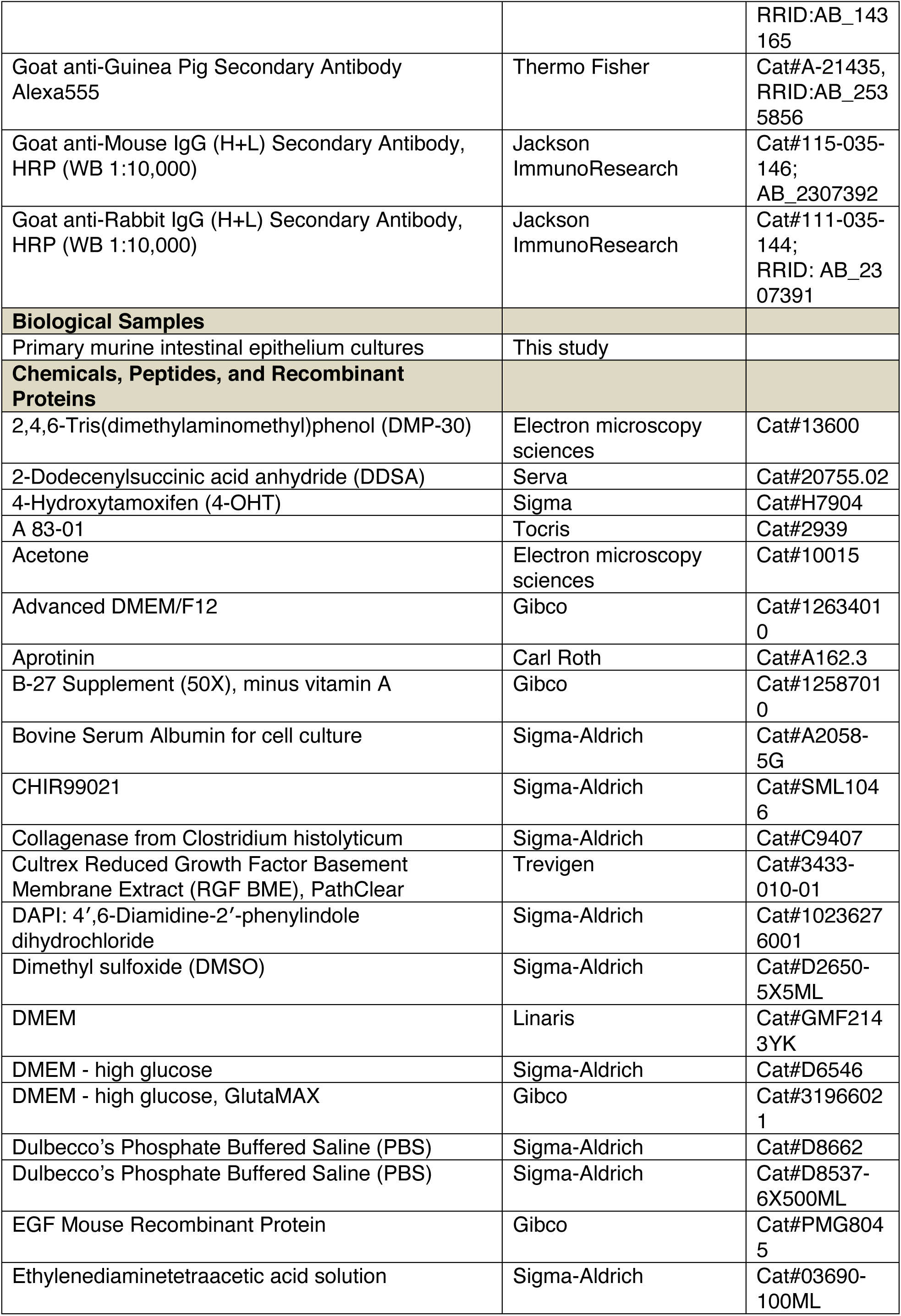
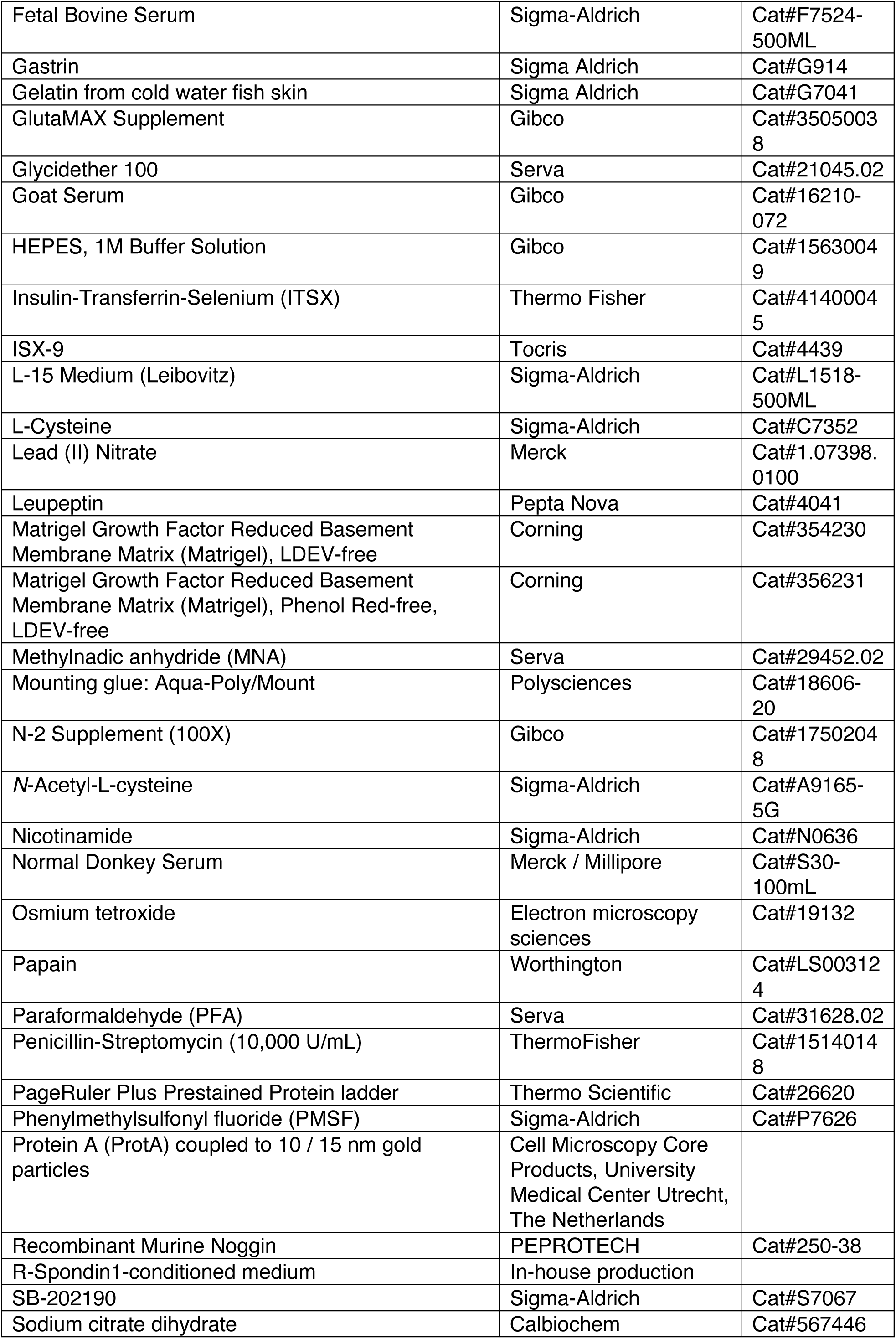
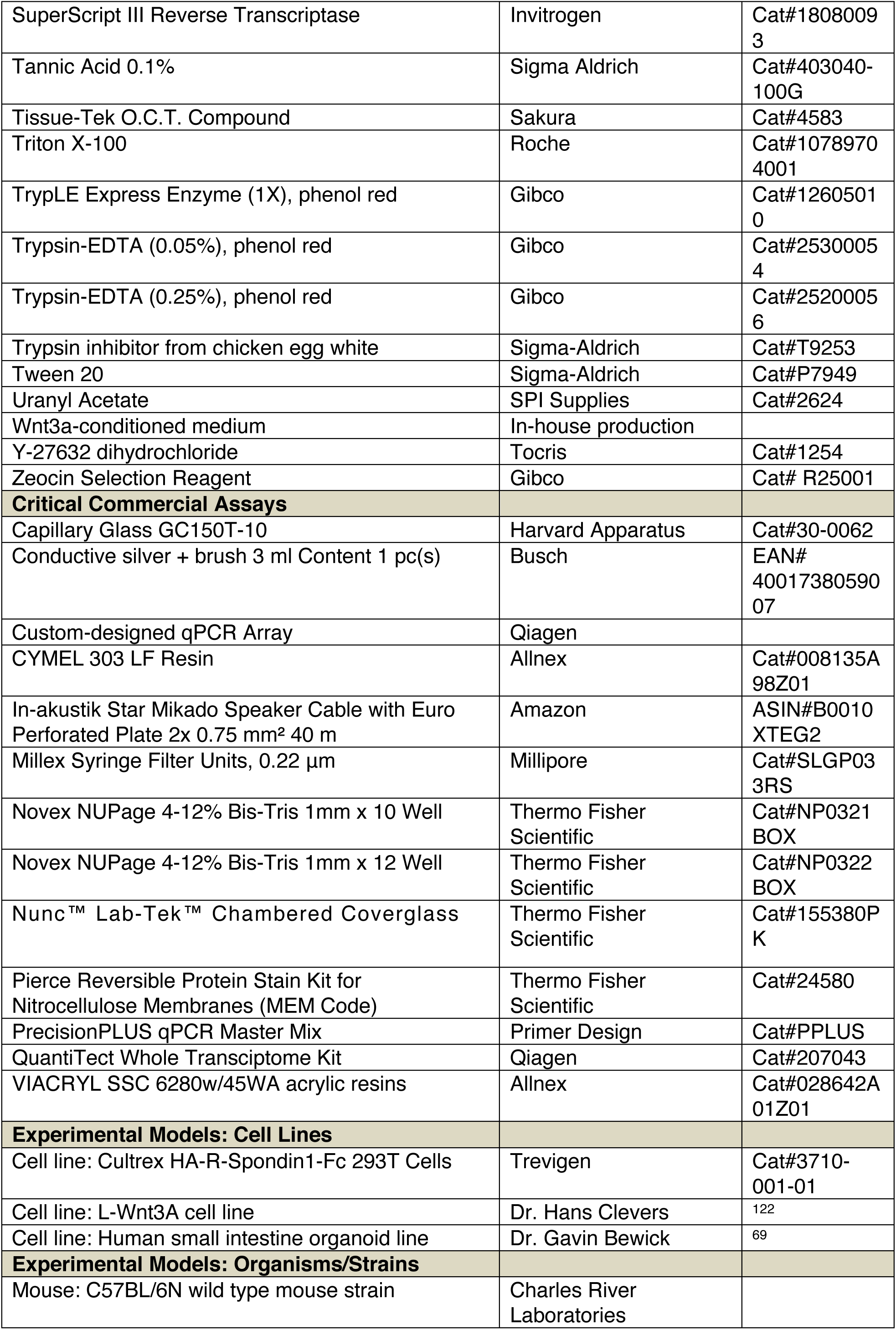
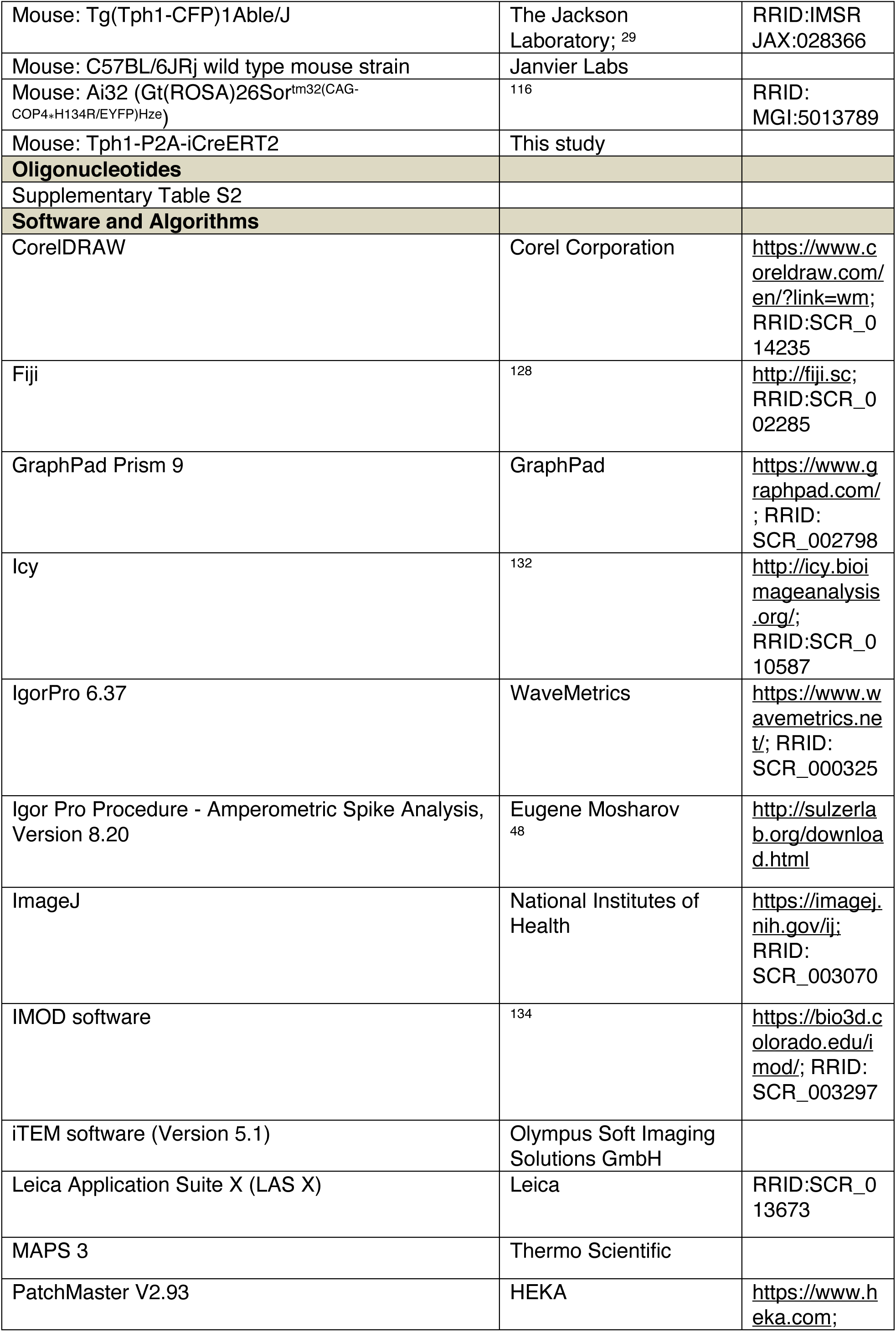
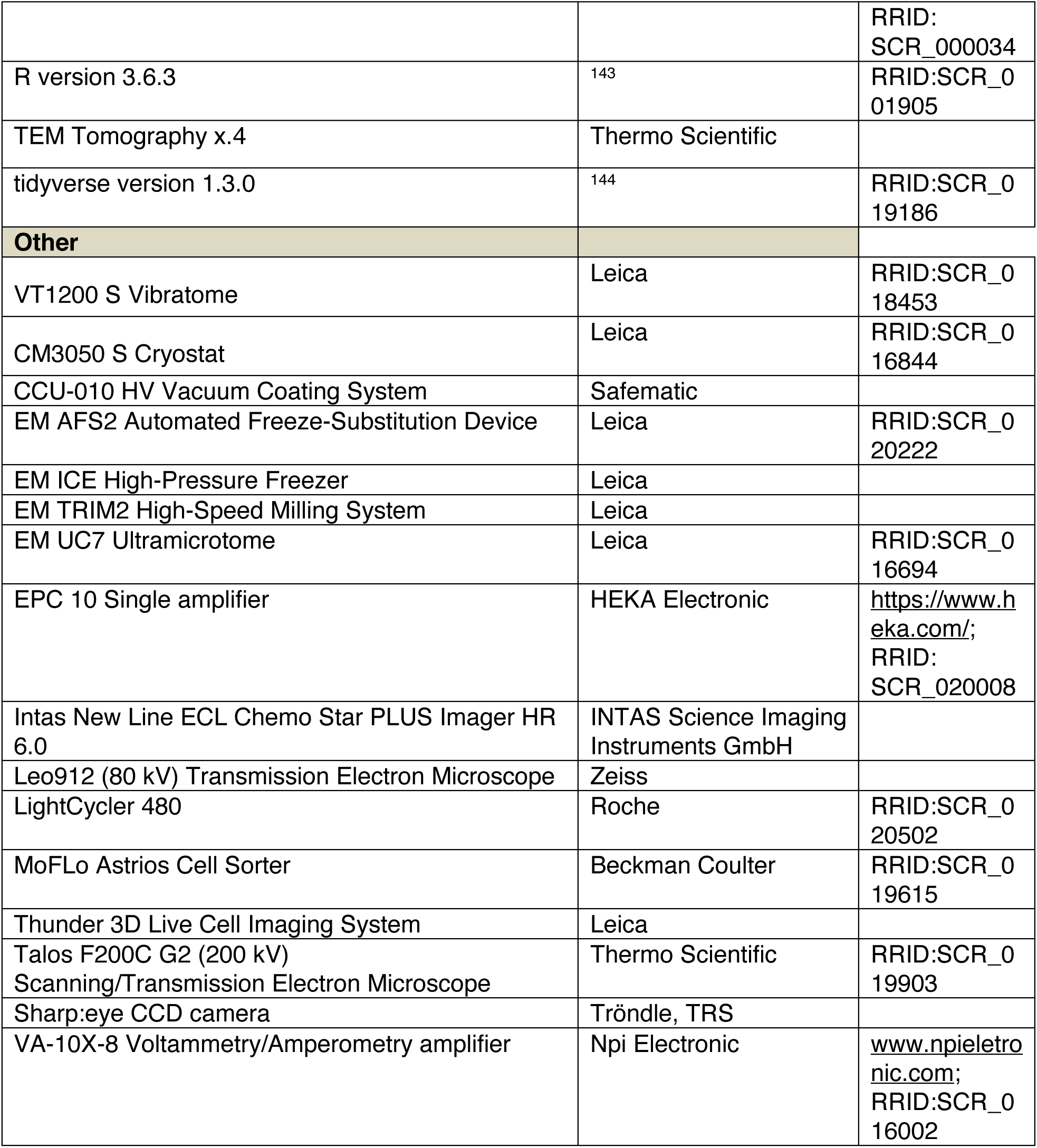

---

**Supplementary Figure S1.**
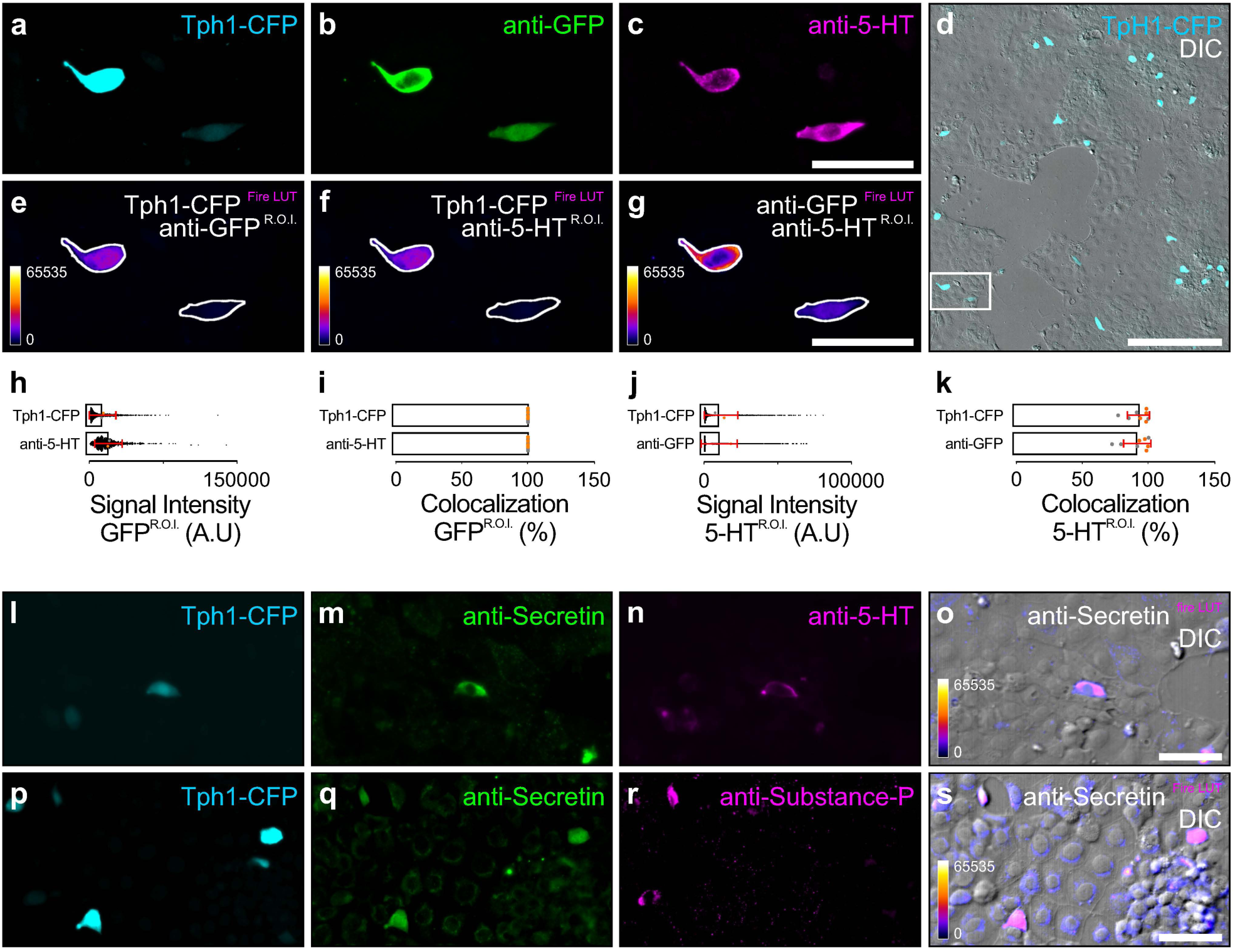
Light Microscopic Analysis of Cultured EC Cells from the Tph1-CFP Mouse Line in Intestinal Epithelial Monolayer Cultures. **a**-**g** Co-labeling of CFP-fluorescent cells (**a** and **d**, cyan) with anti-GFP (**b**, green) and anti-5HT (**c**, magenta) antibodies. The region shown in (**a**-**c**) and (**e**-**g**) is indicated by the insert in **d** (overlay endogenous CFP signal and DIC image). **e**-**f** Co-localization experiments to determine signal intensities of the endogenous CFP fluorescence (fire LUT) in ROIs created in ImageJ based on the anti-GFP signal (**e**), and the anti-5-HT signal channel (**f**). **g** Co-localization experiments to determine signal intensities of the anti-GFP enhanced CFP signal (fire LUT) in ROIs created in ImageJ based on the anti-5-HT signal channel. **h** Intensities of endogenous CFP fluorescence and anti-5HT immunofluorescent signals in ROIs (1118 ROIs analyzed) based on anti-GFP signals. **i** Percent colocalization of endogenous CFP fluorescence and anti-5HT immunofluorescence in ROIs created based on anti-GFP signals. Data points originating from different cultures are separated by color. **j** Intensities of endogenous CFP fluorescence and anti-GFP immunofluorescent signals in ROIs (1930 ROIs analyzed) created based on anti-5-HT signals. **k** Percent colocalization of endogenous CFP fluorescence and anti-GFP immunofluorescence in ROIs created based on anti-5-HT signals. Data points originating from different cultures are separated by color. **l**-**s** Co-labeling of CFP-fluorescent cells (**l** and **p**, cyan) with anti-Secretin (**m** and **q**, green), and anti-5HT (**n**, magenta) or anti-Substance P (**r**, magenta) antibodies. Overlay image showing the anti-Secretin immunofluorescent signal (**o** and **s**, fire LUT) superimposed on the corresponding DIC image (**o** and **s**). Error bars indicate mean ± SD. ORG, 2 cultures, n = 8 coverslips. Scale bars: 50 *µ*m (**c**, **g**, **o**, **s**); 200 *µ*m (**d**).

**Supplementary Figure S2.**
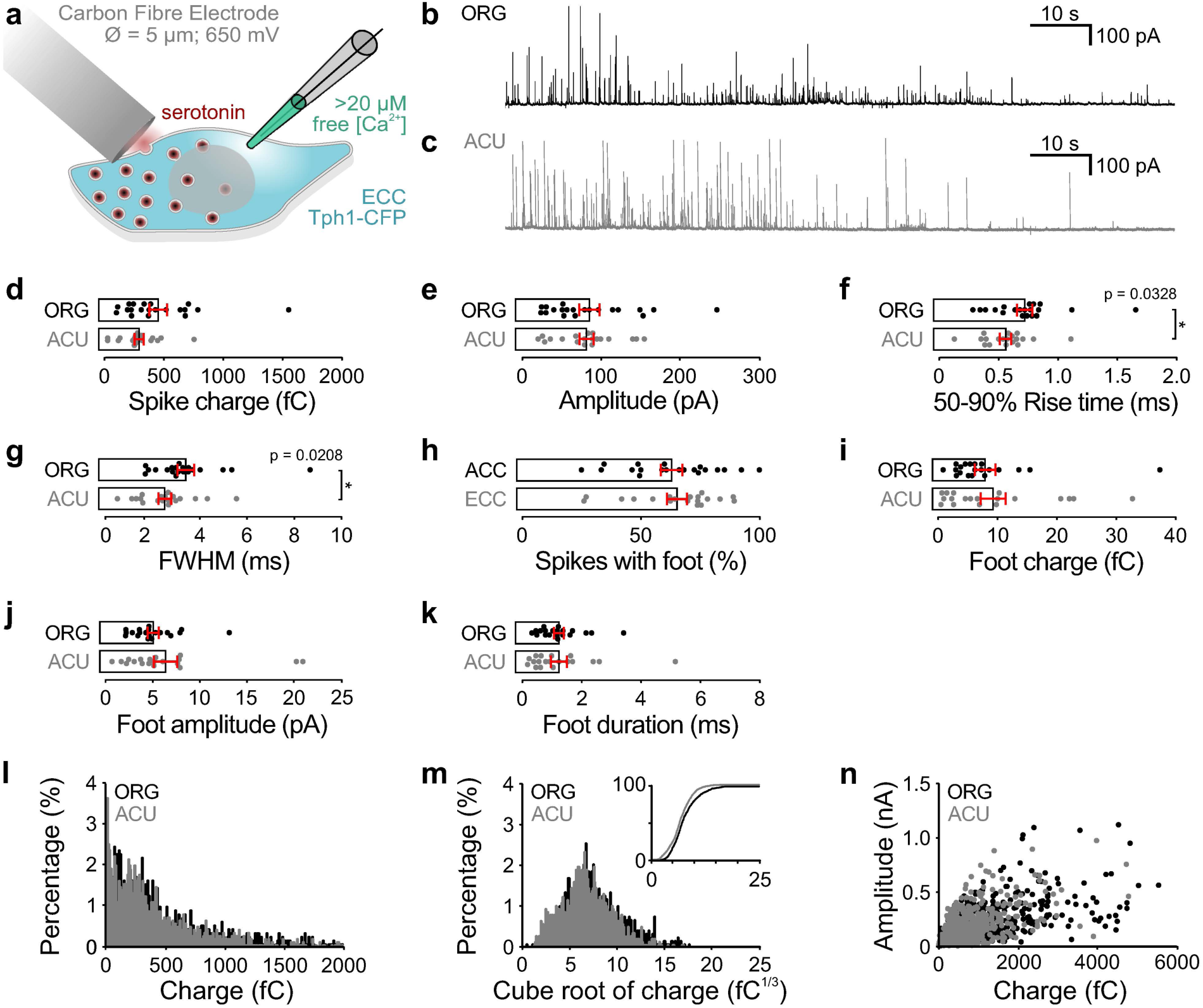
Comparison of Monoamine Release from Mouse EC Cells (ECC) in Acute or Organoid-Derived Epithelial Monolayer Cultures. (**a**) Exocytosis in ECCs was triggered by infusing individual cultured cells with a solution containing high (> 20 *µ*m) [Ca^2+^]. Serotonin release was measured as described in Fig. 3. (**b**-**c**) Representative amperometric recording of an ECC in organoid-derived (ORG) or acutely prepared (ACU) cultures. (**d**) Median spike charge (fC). (**e**) Median maximum spike amplitude (pA). (**f**) Median 50-90% rise time (ms). (**g**) Median full-width half maximum (FWHM) corresponding to the width (ms) at half maximal amplitude. (**h**) Percentage of spikes with a foot signal from all spikes analyzed per cell (%). (**i**) Median foot charge (fC). (**j**) Median maximum foot amplitude (pA). (**k**) Median foot duration (ms). (**l**) Frequency distribution of the amperometric spike charge (fC) for all events measured over all cells [bin width 10 fC, only events <2000 pA; ORG, 1080 spikes, ACU, 1323 spikes]. (**m**) Frequency distribution plotting the cube-root of amperometric spike charges (fC^1/3^) for all events measured over all cells [bin width 0.1 fC^1/3^; ORG, 1163 spikes, ACU, 1350 spikes]. Insert, cumulative frequency distribution for the same data. (**n**) Scatterplot illustrating the relationship between amperometric spike charge (fC) and maximum amplitude (nA) for each event measured [ORG, 1163 spikes, ACU, 1350 spikes]. Error bars indicate mean ± SEM; p < 0.05; **p < 0.01; ***p < 0.001. ORG, 3 cultures, 20 cells; ACU, 5 cultures, 19 cells.

**Supplementary Figure S3.**
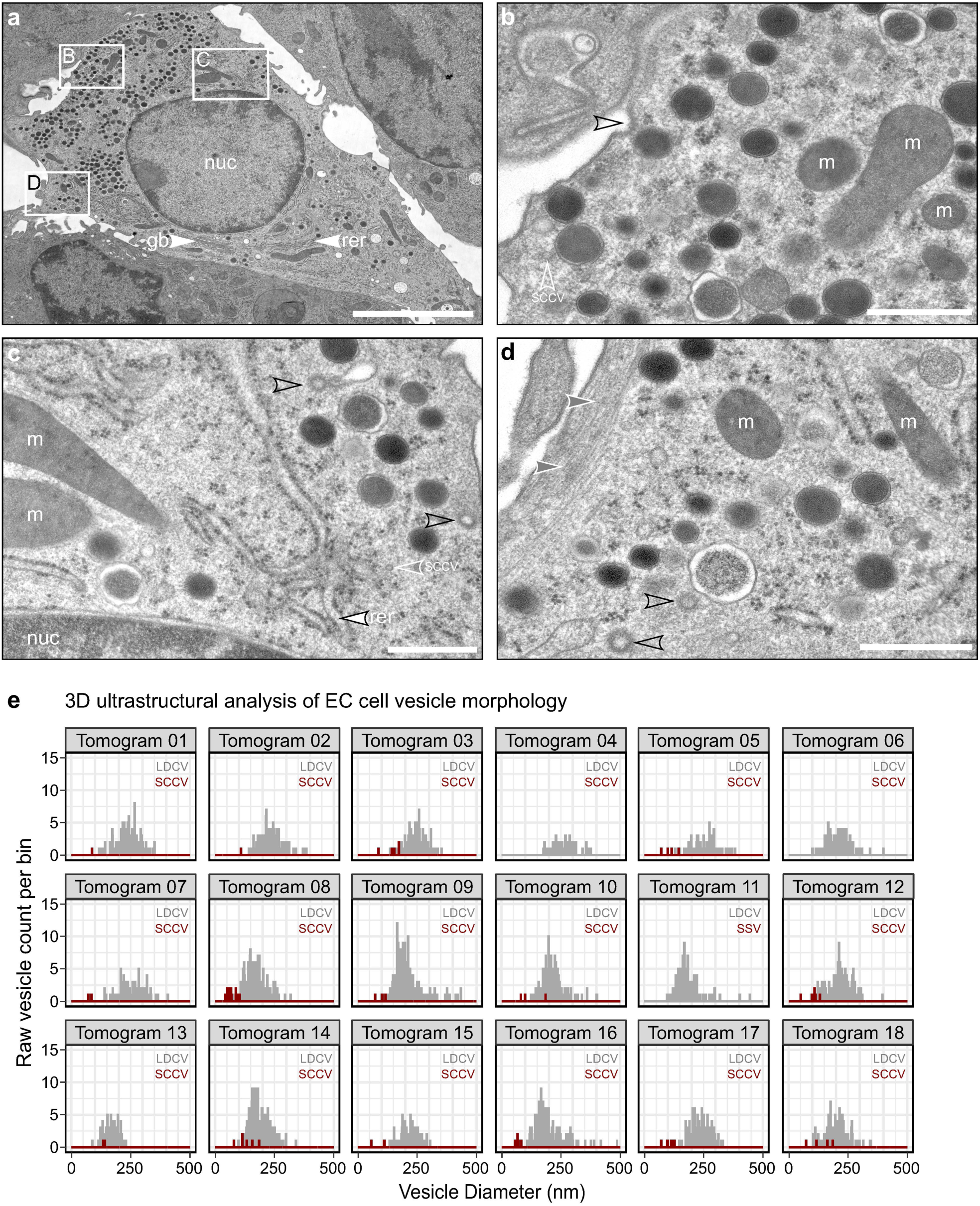
Ultrastructural Properties of Cultures EC Cells. **a**-**d** High magnification electron micrograph through a cross section of the EC cell in position 1 in Fig. 5 for the visualization of ultrastructural features and organelles [black open arrowheads, coated membrane pits and vesicles; grey filled arrowheads, submembranous filament bundles; white arrowheads, rough endoplasmic reticulum (rer) and golgi body (gb); white open arrowheads, small clear core vesicle (SCCV); m, mitochondria; nuc, nucleus]. **e** Distribution of vesicle diameters separated by SCCVs and LDCVs for the individual tomograms analyzed [bin width 2 nm] (corresponding to pooled data in Fig. 6). Scale bars: 5 μm (**a**); 500 nm (**b**-**d**).

**Supplementary Figure S4.**
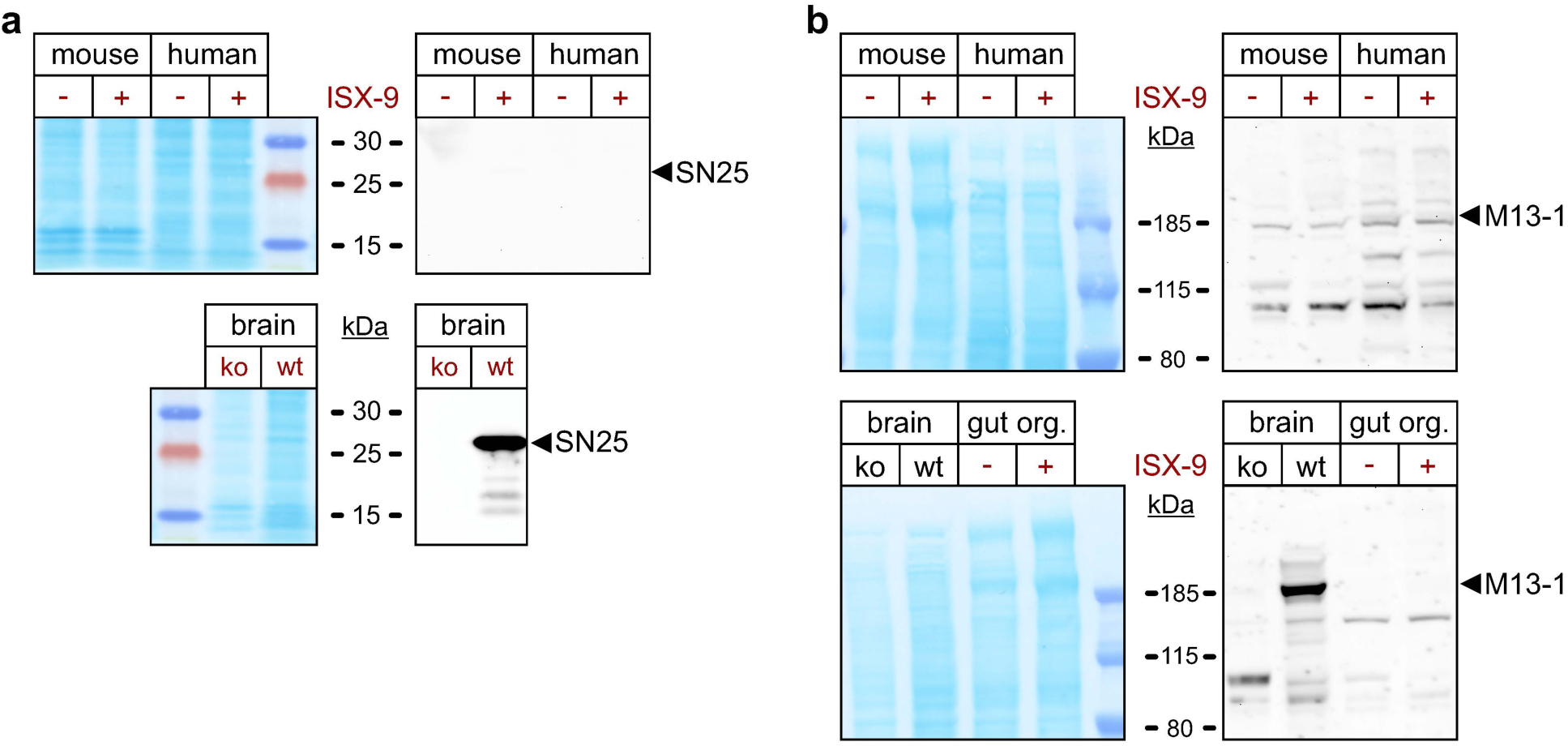
Characterization of the Expression of Candidate Components of the Presynaptic Neurotransmitter Release Machinery in EC cells by Protein Immunoblotting. **a** Western blot analysis of mouse and human organoids treated with ISX-9 (+) or DMSO (-) using an antibody directed against SNAP-25 (SN25, top panel). Protein stain of the nitrocellulose membrane (MEM Code, left panel) and chemiluminescent signals detected for the same membrane segments after immunolabelling with different antibodies (right panels). No chemiluminescent signals were detected with the antibody in intestinal organoid homogenates, while it produced a highly specific signal in wildtype (wt) brain homogenate samples that was absent from SNAP-25 knockout (ko) brain homogenates (Lower panel). **b** Western blot analysis of mouse and human organoids treated with ISX-9 (+) or DMSO (-) using anti-Munc13-1 antiserum (M13-1, top panel). No specific immunoreactive bands were detected with the antiserum in intestinal organoid homogenates at the expected size of the protein. The same antiserum produced a highly specific signal in wildtype (wt) brain homogenate samples, which was absent in Munc13-1 and -2 knockout (ko) brain homogenates (Lower panel).

**Supplementary Figure S5.**
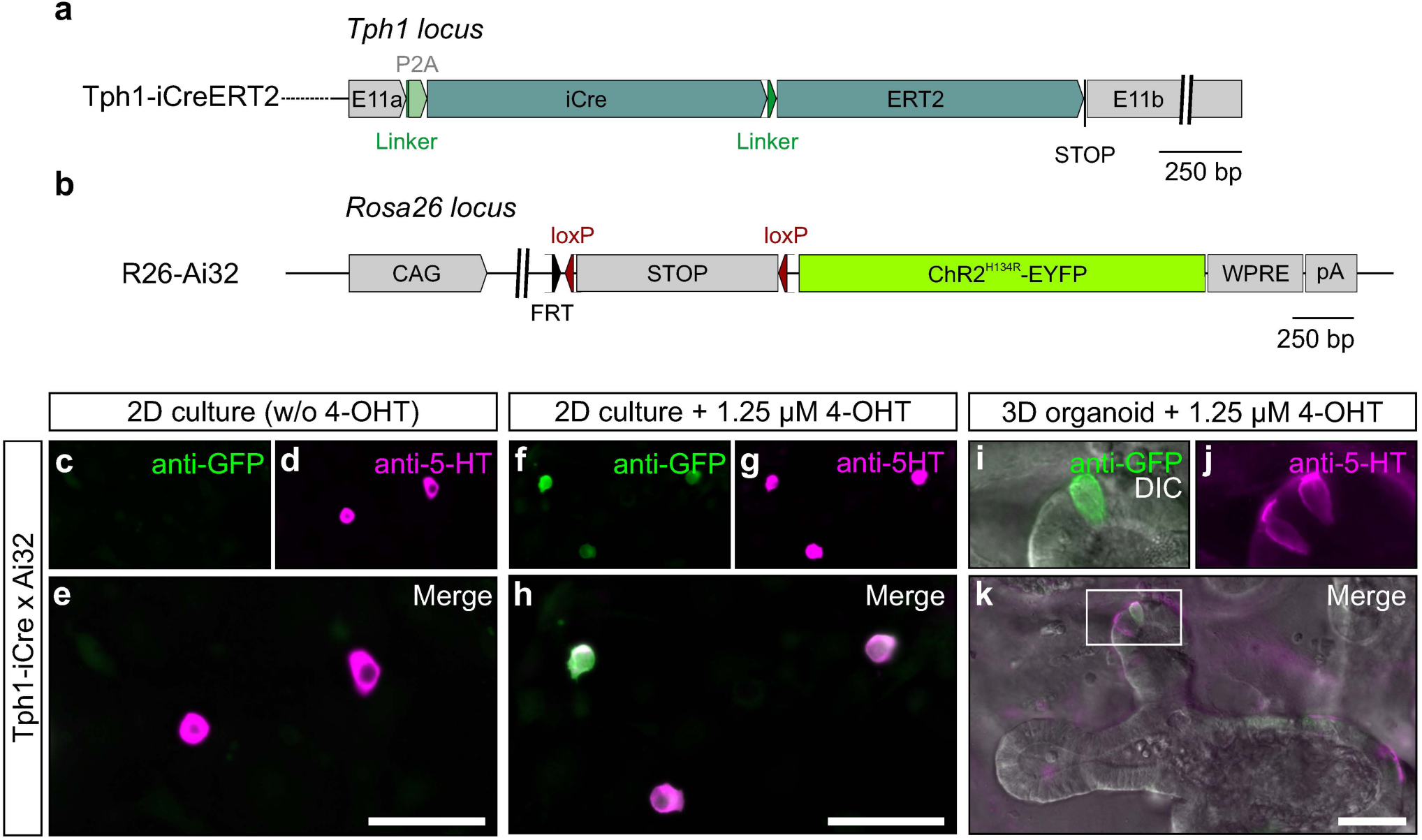
Generation and Validation of a New Tph1-iCreERT2 Line for the Cell-Specific and Temporally Controlled Expression of Fluorescent Reporters and Optogenetic Actuators in Cultured EC Cells. **a** Illustration of the Tph1-P2A-iCreERT2 knockin (KI) allele (P2A, self-cleaving peptide; iCre, codon-improved Cre-recombinase; ERT2, estrogen receptor targeting motif; STOP, Stop codon) generated in this study. **b** Illustration of the Ai32 KI allele^116^ in the Rosa26 (R26) locus (CAG, CAG promoter; FRT-loxP-STOP-loxP, loxP-flanked STOP cassette, ChR2. Channelrhodopsin 2; EYFP, enhanced green fluorescent protein; WPRE, Woodchuck Hepatitis Virus Posttranscriptional Regulatory Element; pA, poly-A tail). **c**-**h** Immunofluorescence images of untreated (**c**-**e**) and 4-hydroxytamoxifen (4-OHT) treated (**f**-**h**) Tph1-iCreERT2;Ai32 colonic epithelial monolayer cultures stained with antibodies directed against GFP (green, **c**, **e**, **f**, **h**) and 5-HT (magenta, **d**, **e**, **g**, **h**). Treatment of cultures with a 16 h pulse of 4-OHT induced prominent expression of the ChR2-EYFP fusion protein in 5-HT positive cells (**f**, **h**). **i**-**k** Immunofluorescence images of 4-OHT treated Tph1-iCreERT2;Ai32 duodenal organoid. Translocation of iCreERT2 to the nucleus was triggered with 1.25 *µ*m 4-OHT on DIV4 for 15 h and organoids were fixed and processed for immunofluorescence imaging 5 days later. Using this protocol, only a subset of 5-HT immunopositive cells (magenta, **j** and **k**) demonstrate expression of the ChR2-EYFP fusion protein (green, **I**, **k**). These data indicate that it is possible to temporally control the expression of reporters in EC cells. Scale bars: 50 *µ*m (**e**, **h**, **l**).

